# FTO-mediated cytoplasmic m^6^A_m_ demethylation adjusts stem-like properties in colorectal cancer cell

**DOI:** 10.1101/2020.01.09.899724

**Authors:** Sébastien Relier, Julie Ripoll, Hélène Guillorit, Amandine Amalric, Florence Boissière, Jérôme Vialaret, Aurore Attina, Françoise Debart, Armelle Choquet, Françoise Macari, Emmanuelle Samalin, Jean-Jacques Vasseur, Julie Pannequin, Evelyne Crapez, Christophe Hirtz, Eric Rivals, Amandine Bastide, Alexandre David

**Author notes:** Co-last and corresponding authors (E.R.), (A.B.) and (A.D.).

## Abstract

Cancer stem cells (CSCs) are a small but critical cell population for cancer biology since they display inherent resistance to standard therapies and give rise to metastases. Despite accruing evidence establishing a link between deregulation of epitranscriptome-related players and tumorigenic process, the role of messenger RNA (mRNA) modifications dynamic in the regulation of CSC properties remains poorly understood. Here, we show that the cytoplasmic pool of fat mass and obesity-associated protein (FTO) impedes CSC abilities in colorectal cancer through its m^6^A_m_ (N^6^,2’-O-dimethyladenosine) demethylase activity. While m^6^A_m_ is strategically located next to the m^7^G-mRNA cap, its biological function is not well understood and has not been addressed in cancer. Low FTO expression in patient-derived cell lines elevates m^6^A_m_ level in mRNA which results in enhanced *in vivo* tumorigenicity and chemoresistance. Inhibition of the nuclear m^6^A_m_ methyltransferase, PCIF1/CAPAM, partially reverses this phenotype. FTO-mediated regulation of m^6^A_m_ marking constitutes a novel, reversible pathway controlling CSC abilities that does not involve transcriptome remodeling, but could fine-tune translation efficiency of selected m^6^A_m_ marked transcripts. Altogether, our findings bring to light the first biological function of the m^6^A_m_ modification and its potential adverse consequences for colorectal cancer management.

---

Despite significant advances in diagnosis and therapy, colorectal cancer (CRC) remains a major cause of mortality and morbidity worldwide. CRC survival is highly dependent upon early diagnosis. Patients with localized cancer exhibit 70 to 90% 5-year survival. Survival from metastatic cancer plummets to 10%. Metastasis is a multistep process encompassing local infiltration of tumor cells into adjacent tissues, transendothelial migration into vessels, survival in the circulatory system, extravasation, and colonization of secondary organs [1]. This process entails constant reprogramming of gene expression to enable tumor adaptation in different environments, a peculiar trait of cancer stem cells (CSCs). CSC constitute a minor subpopulation of tumor cells endowed with self-renewal and multi-lineage differentiation capacity [2]. The most clinically relevant trait of CSCs is their ability to metastasize and escape from standard chemotherapy [3]. Understanding the molecular mechanisms that participate to the CSC phenotype is critical to designing improved cancer therapeutics.

Among more than 100 post-transcriptional modifications reported to occur on RNA [4], N6-methyladenosine (m^6^A) is the most frequent epigenetic modification of mammalian messenger RNAs (mRNAs) [5]. m^6^A is involved in all post-transcriptional steps of gene expression (mRNA splicing, transport, stability and translation) and plays a role in pleiotropic biological processes including development, immunology, and stem cell biology [5]. Therefore, it comes at no surprise that m^6^A dysregulation is intricately involved in the progression of several solid and non-solid tumors while the underlying functional mechanism differs from one malignancy to another [6-11].

Discovered several decades ago [12-16], the function of m^6^A remained obscure until the identification of the first m^6^A demethylase, the fat mass and obesity-associated protein (FTO) [17]. The marriage of immunochemical approaches with next-generation sequencing (NGS) technologies revealed the unique topology of m^6^A distribution along mRNA. m^6^A is a dynamic reversible chemical modification catalyzed by a protein complex consisting of the methyltransferase-like 3 and 14 (METTL3 and METTL14), and several auxiliary proteins such as the Wilms’ tumor 1-associating protein (WTAP) [18-21]. Effects of m^6^A involve recruitment of reader proteins, e.g. YTHDF1 [22] or YTHDF2 [23], that lodge the modified adenosine in the hydrophobic pocket of their YTH domain. Finally, m^6^A is removed by the AlkB homolog 5 (ALKBH5) [24] and FTO [17] (**Figure 1a**). m^6^A modification generally occurs in a subset of RRA*CH consensus sites (R, purine; A*, methylable A; C, cytosine; H, non-guanine base), at the beginning of the 3′-UTR near the translation termination codon [25, 26] with one exception. Indeed, besides internal adenosine, 2′-O-methyladenosine (A_m_) residue adjacent to the N7-methylguanosine (m^7^G) cap, can be further methylated at the N6 position and become the N6,2′-O-dimethyladenosine (m^6^A_m_) [27]. m^6^A_m_ can be deposited by the recently identified PCIF1/CAPAM [28] and removed by FTO [29]. Unlike m^6^A, the biological function of m^6^A_m_ mRNA modification is poorly understood and its potential involvement in cancer onset or evolution has never been addressed.

**Figure 1.**
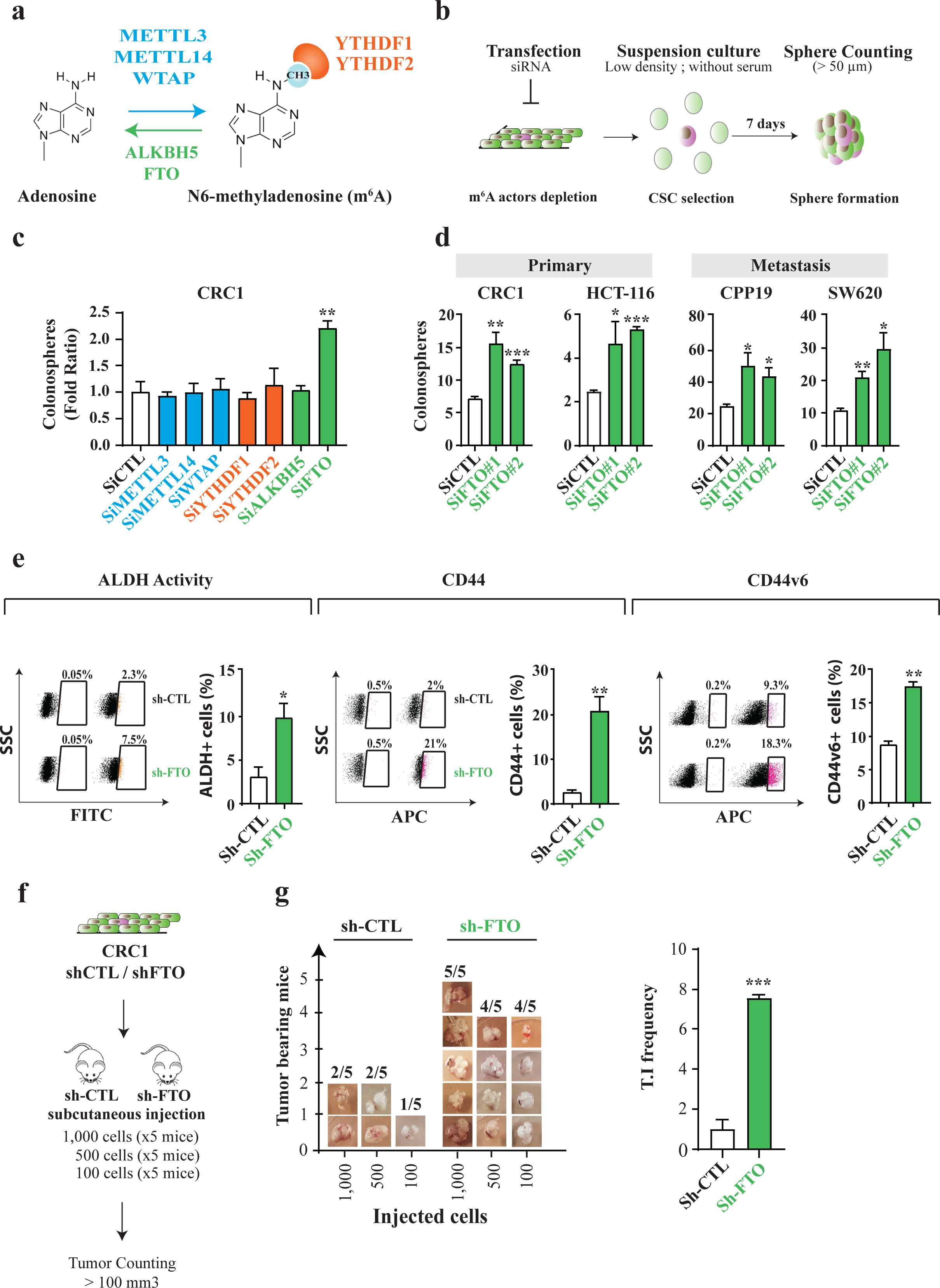
FTO inhibition promotes stem-like traits in colorectal cancer cell lines. **(a) Actors of m**^**6**^**A modification targeted by the siRNA screening**. The writer complex METTL3 – METTL14 – WTAP deposits m^6^A while ALKBH5 and FTO erases m^6^A. Both YTHDF1 and YTHDF2 are readers of m^6^A modification. **(b) Sphere forming assay**. siRNA-transfected cells were seeded at low density (1 cell/µL) in non-adherent conditions and serum-deprived medium. This type of suspension culture allows the survival and growth of stem-like/progenitor cells. Following 7 days of culture, the number of spheres correlates with the initial number of CSC. **(c) FTO silencing increases the sphere forming potential of CRC1 cell line**. Colonosphere formation was quantified following knockdown of the main m^6^A actors. Results are expressed in fold change compared to si-CTL. n = 3 biological replicates. Mean +/- S.E.M, ** p-value < 0.01. Two-sided Unpaired T-test. **(d) FTO silencing increases the sphere forming potential in four CRC cell lines**. Colonospheres quantification after silencing of FTO by two distinct siRNA, in four cell lines derived from primary tumors or metastasis. Results are expressed in fold change compared to si-CTL (n = 3 biological replicates). Mean +/- S.E.M, * p-value < 0.05; ** p-value < 0.01. Two-sided Unpaired T-test. **(e-g) Expression of CSC biomarkers**. Level of ALDH activity **(e)** as well as cell-surface expression of CD44 **(f)** and CD44v6 **(g)** were evaluated by flow cytometry in CRC1-sh-FTO cell line *vs* sh-CTL. Graphs show one representative biological replicate (n = 3). Bar plots show quantification of the number of ALDH **(e)**, CD44 and CD44v6 positive cells from three biological replicates. Mean +/- S.E.M, *p-value < 0.05, **p-value < 0.01. Two-sided Unpaired T-test. **(f) *In vivo* tumor initiation assay**. Either 1,000, 500 or 100 CRC1 sh-FTO / sh-CTL cells were subcutaneously injected into nude mice. After 7 weeks, the number of tumor-bearing mice (tumor > 100 mm^3^) was evaluated. **(g) FTO silencing increases tumor initiation *in vivo***. Picture represents the number of tumor bearing mice for each group (5 mice per group) after injection of sh-FTO or sh-CTL CRC1 cell line. Bar plot represents the quantification of Tumor Initiation (T.I) frequency obtained by ELDA software. *** p-value < 0.001, chi2 test.

Here, we uncover the importance of cytoplasmic FTO-mediated m^6^A_m_ level adjustment to colorectal CSC. Using patient-derived cell lines, we establish that FTO activity hampers CSC abilities through a novel, unconventional process that does not involve transcriptional reprogramming but, most likely, fine-tuning translation of m^6^A_m_ marked transcripts.

## RESULTS

### FTO inhibition promotes stem-like traits in colorectal cancer cell lines

We initially evaluated the involvement of m^6^A modification in generating the CSC phenotype. Due to their inherent plasticity, CSCs are best identified via their functional abilities, such as tumorigenic potential and chemoresistance, rather than surface biomarkers. We examined the ability of short interfering RNAs (siRNA) targeting known m^6^A mediators –writers, readers and erasers-to alter sphere-forming potential (SFP) (**Figure 1b**). SFP is the ability of cancer cells-from either conventional cell lines or patient exeresis-to form microtumor-like spheroids (colonospheres) from a single cancer progenitor cell [30]. This model is often used as a surrogate to evaluate the tumorigenic potential of solid tumors (5). We used CRC1 cells, a colorectal cancer cell line established in the lab from colorectal cancer exeresis [31]. The various siRNAs significantly silenced expression of individual target genes: METTL3, METTL14, WTAP, YTHDF1, YTHDF2, ALKBH5 and FTO (**Figure S1a**). Only FTO knockdown affected SFP, nearly doubling colonospheres numbers (**Figure 1c)**. We confirmed the importance of FTO using a different targeting siRNA (**Figure 1d, S1b**) and three other cell lines, derived from primary (HCT-116) and metastatic (CPP19, SW620) tumors (**Figure 1d**).

Next, we generated stable CRC1 cell lines expressing GFP with either a short hairpin RNA targeting FTO (sh-FTO) or an irrelevant scrambled short hairpin control (sh-CTL), and selected transfected clones by cell sorting for GFP expression (**Figure S1c**). Sh-FTO expressing cell lines established from individual cells exhibited a three to ten-fold decrease of FTO expression as shown by immunoblot analysis (**Figure S1d**) and displayed increased SFP, confirming the siRNA findings (**Figure S1e**). Importantly, FTO knockdown does not influence cell growth (**Figure S1f**), which differs from the AML (acute myeloid leukemia) model [8]. Then, we evaluated the level of several stemness-related markers associated with CSC features. FTO knockdown cells displayed enhanced aldehyde dehydrogenase (ALDH) activity, a hallmark of CSC [32] (**Figure 1e**), and increased expression of CD44 and CD44v6 (**Figure 1e**), membrane receptors associated with tumor progression [33] and indispensable for CSC tumor initiation, chemoresistance, and epithelial to mesenchymal transition. CRC1 sh-FTO cells demonstrated a ten-fold increase in the number of CD44+ cells (from 2.1% to 21%) (**Figure 1e**) and CD44v6 isoform expression was doubled (**Figure 1e**). FTO knockdown in SW620 cells phenocopied the CRC1 cells (**Figure S1c-h**).

To connect FTO levels with *in vivo* tumor initiation potential we inoculated immunodeficient mice (athymic nude) with increasing numbers of sh-FTO or sh-CTL cells (**Figure 1f**). Seven weeks later, tumor xenografts (diameter > 100mm^3^) were counted and harvested. Remarkably, as few as hundred sh-FTO cells were capable of initiating tumor formation in 4/5 mice *vs*. 1/5 for sh-CTL cells (**Figure 1g**). Extreme limiting dilution software analysis (ELDA) shows that the frequency of tumor-initiating cells is seven-fold higher in sh-FTO cells, with a highly significant p-value (p=0.00035) (**Figure 1g**).

Based on these findings, we conclude that diminished FTO expression promotes the CSC phenotype.

### FTO silencing confers resistance to chemotherapy in colorectal cancer cell lines

We extended these findings to chemoresistance, another hallmark of CSC. We employed FIRI treatment, a combination of 5-fluorouracil and irinotecan, used to treat metastatic colorectal cancer [34]. We followed a standard protocol for treating colorectal cancer cell lines based on 3 days treatment with FIRI (50μM 5-flurouracil (5-FU) + 500nM SN38, active metabolite of irinotecan) (**Figure 2a**). siRNA mediated FTO targeting conferred significant chemoprotective effects on CRC1 cells (2 to 3-fold) in comparison with control cells (si-CTL)(**Figure 2a**). As above, we obtained a similar effect in SW620 (**Figure 2a**), as well as in stable sh-FTO models (**Figure 2b**).

**Figure 2.**
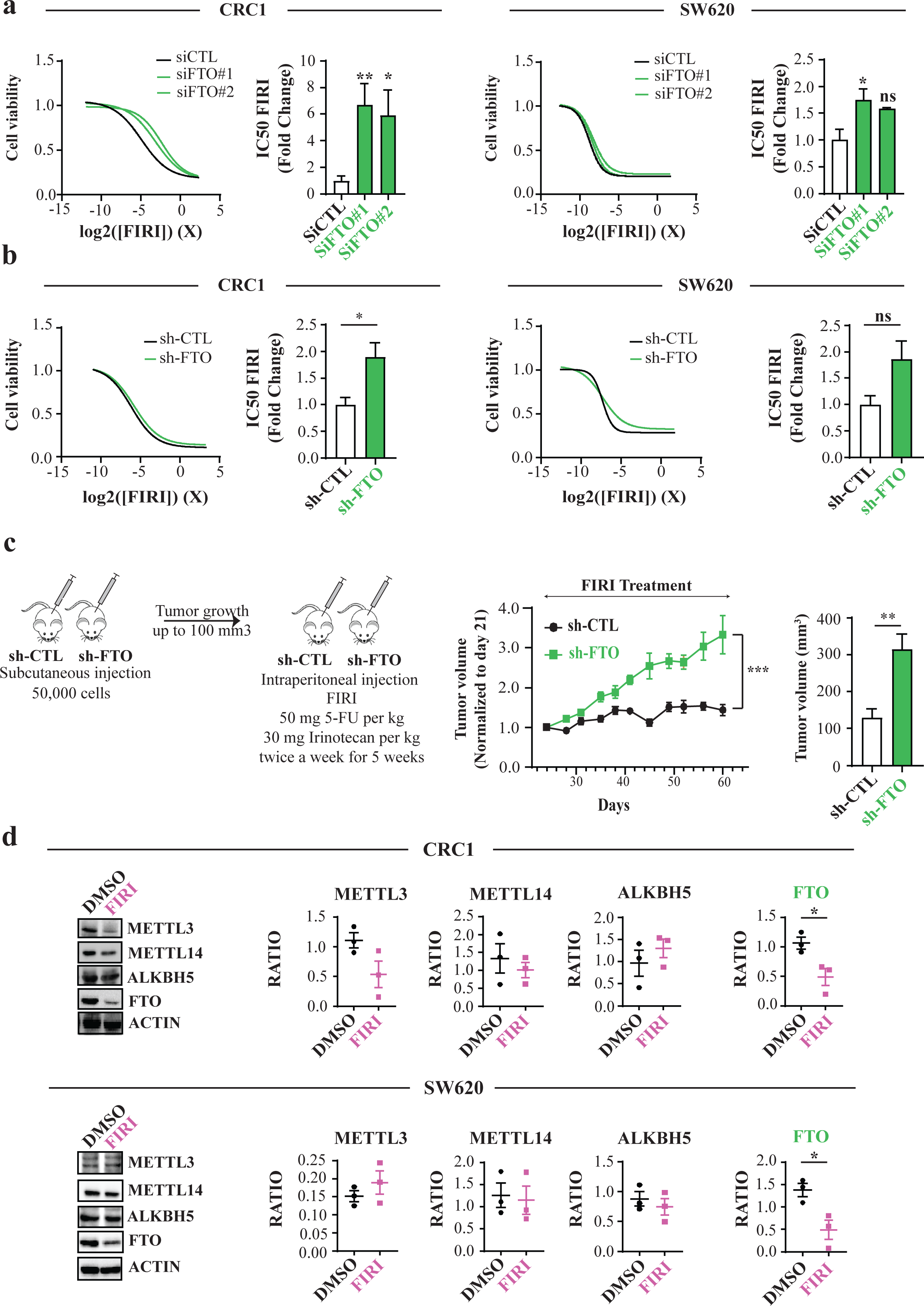
FTO silencing confers resistance to chemotherapy. **(a) Transient FTO silencing increases chemoresistance to FIRI.** Graphs illustration of FIRI toxicity on either si-FTO or si-CTL-transfected cells (CRC1 (left) and SW620 (right) cell lines). Toxicity was measured using Sulforhodamine B assay. Bar plot represents quantification of IC_50_ of three biological replicates. FIRI 1 X = 50 µM 5-FU, 0.5 µM SN38. Mean +/- S.E.M, **p-value < 0.01,*p-value < 0.05, Two-sided Unpaired T-test. **(b) Stable FTO silencing increases chemoresistance to FIRI**. Same as **(a)** with stable cell lines (sh-FTO or sh-CTL) from two distinct backgrounds (CRC1 (left) and SW620 (right)). Mean +/- S.E.M, *p-value < 0.05, Two-sided Unpaired T-test. **(c) FTO silencing increases *in vivo* chemoresistance**. 50,000 of either sh-FTO or sh-CTL cells were subcutaneously injected into the flank of nude mice. After 21 days (tumor size about 100mm3), mice were treated with FIRI (50 mg FIRI per kg, 30 mg irinotecan per kg) and tumor growth was measured twice a week for 5 weeks. *** p-value < 0.001, two-way ANOVA test. Bar plot represent mean +/- S.E.M of tumor volume measured at the last time point. **p-value <0.01. **(d) FIRI-resistant cells display decreased FTO expression**. Immunoblot analysis of METTL3 – METTL14, ALKBH5 and FTO levels after 72 h of 0.2 X FIRI treatment. Pictures are representative of 3 experiments in CRC1 cell line. Protein level quantification is mean +/- S.E.M of FTO normalized to ACTIN of three biological replicates. * p-value < 0.05, ns = not significant, Two-sided Unpaired T-test.

Importantly, chemoprotection bestowed by FTO silencing extended to FOX treatment (50μM 5-flurouracil (5-FU) + 1 µM oxaliplatine), another advanced stage colorectal cancer therapy [35, 36] (**Figure S2a**). To evaluate the effect of FTO knockdown *in vivo* we injected sh-CTL or sh-FTO cells (50,000 cells) into nude mice (6 mice per group). When the tumors reached about 100 mm^3^ (day 21), we treated mice with FIRI (50mg/kg 5-FU + 30mg/kg irinotecan) and measured tumors twice a week for four and a half weeks (**Figure 2c**). While FIRI treatment stabilized tumor size in sh-CTL mice, sh-FTO tumors displayed substantial chemoresistance maintaining consistent growth (**Figure 2c**).

Do chemoresistant cells display low FTO levels? To produce CSC-enriched chemoresistant cells, we treated CRC1 and SW620 cells for three days with sub-lethal doses of FIRI (0.2 X FIRI = 10 µM 5-FU + 0.1µM SN38). As expected, 0.2 X FIRI treatment triggered cell cycle arrest at G2/M phase and increased subG1 cells (**Figure S2b**). Surviving cells demonstrated increased ALDH activity (**Figure S2c**) and enhanced resistance to chemotherapy (**Figure S2d**). While variations of METTL3 and ALKBH5 levels could be noticed following chemoresistance acquisition, they were not consistent between cell lines (**Figure 2d**). By contrast, FTO level was decreased by half in both cell lines (**Figure 2d**). Noteworthy, the correlation between FTO mRNA expression levels and protein abundance varies from one cell line to another (**Figure S2e**).

Together, these findings demonstrate that FTO expression is tightly linked to the CSC chemoresistant phenotype.

### FTO expression regulates CSC phenotype in circulating tumor cells

Circulating tumor cells (CTCs) are often detected in the bloodstream of colorectal cancer patients, sometimes even at early stages of the disease [37]. CTC are responsible for metastasis and high mortality. We previously established CTC lines from chemotherapy-naïve stage IV (metastatic) CRC patients that display a strong CSC phenotype [31]. Immunoblotting revealed that FTO protein levels are reduced by ∼50% in CTC lines in comparison with primary and metastatic patient derived lines (**Figure 3a**). Decreased expression is achieved post transcriptionally, since quantitative PCR (qPCR) analysis of FTO mRNA levels showed no significant differences between CTC lines and primary/metastatic patient derived lines (**Figure 3b**). A similar disconnect in FTO mRNA/protein was reported in gastric tumors [38].

**Figure 3.**
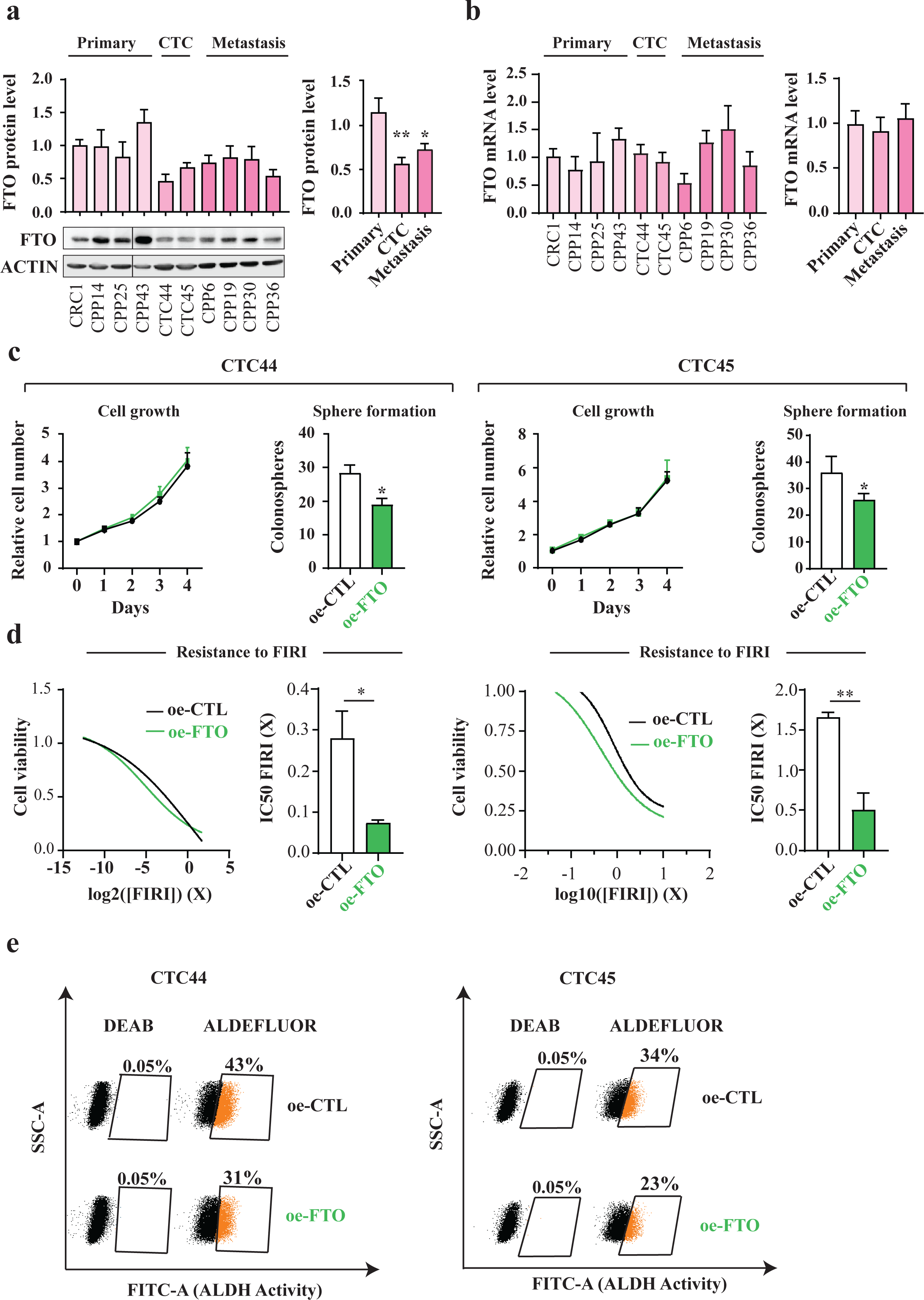
FTO expression is lower in “circulating tumor cells” (CTC) than in “solid” tumor cells. **(a) FTO protein level is the lowest in Circulating Tumor Cell lines.** FTO protein level quantification by western blot in several patient derived cell lines from distinct origin: primary tumor, blood, and metastasis. The first graph represents the level of FTO protein in each individual cell lines. The following bar plot represents the quantification by groups. Mean +/- S.E.M of three biological replicates, ** p-value < 0.01, * p-value < 0.05, One-way Anova followed by multiple comparison. **(b) FTO mRNA level barely fluctuates throughout colon cancer progression**. Level of FTO mRNA was evaluated by RT-PCR in the same cell lines as in **(a)**. Left graph represents of FTO level in each individual cell line. Right bar plot represents the quantification by groups. Mean +/- S.E.M of 3 biological replicates, ns = not significant, One way Anova. **(c) Increase of FTO level decreases the sphere forming potential of CTC cell lines**. Cell growth and sphere forming potential evaluation after overexpression of FTO in CTC44 and CTC45 lines. **(d) Increase of FTO level sensitizes CTC cell lines to chemotherapy**. FIRI toxicity assay on CTC44 or CTC45 after overexpression of FTO. Mean +/- S.E.M, ** p-value < 0.01, *p-value < 0.05, Two-sided Unpaired T-test. **(e) FTO overexpression decreases ALDH activity in CTC cell lines**. Flow cytometry quantification of ALDH positive cells after FTO overexpression. Graphs show the number of ALDH positive cells in one representative biological replicate out of three for CTC44 and CTC45.

Despite this, we could increase FTO protein by transfecting cells with a plasmid containing a FTO cDNA under a strong promoter. Increasing FTO in CTC lines (CTC44 and CTC45) did not affect cell proliferation in monolayer culture but decreased SFP (**Figure 3c**), chemoresistance to FIRI treatment (**Figure 3d**) and ALDH activity (**Figure 3e**) in both cell lines.

Thus, multiple lines of evidence point to the conclusion that FTO is a key factor in maintaining the CSC phenotype.

### FTO functions via its m^6^A_m_ demethylase activity

How does FTO modulate CSC functions? First, we evaluated whether its catalytic activity was essential to achieve this phenotypic outcome. As FTO can demethylate m^6^A and m^6^A_m_ [39-41], we measured their levels in RNAse digested polyadenylated mRNA from both sh-CTL and sh-FTO cells using high-performance liquid chromatography-coupled to tandem mass spectrometry (LC-MS/MS) analysis. At first, we isolated mRNA from cell pellets and verified the purity of our sample preparation by LC-MS/MS (**Figure S3a**) and qPCR (**Figure S3b**). To detect m^6^A_m_ proximal to m^7^G-Cap, we used a previously published method based on including a decapping step (RNA 5’ Pyrophosphohydrolase, RppH treatment) prior to Nuclease P1 treatment [41]. We calibrated the assay using standard curves created with synthetic nucleoside standards (**Figure S3c**).

In both CRC1 and SW620 lines, FTO knockdown did not impact internal m^6^A / A level (**Figure 4a**). However, we clearly observed a significant increase of m^6^A_m_ / A ratio (**Figure 4a**). We performed controls to strengthen our observation: first, FTO silencing affected neither m^6^A / A nor m^6^A_m_ / A ratio in small RNA species, another potential target of FTO (**Figure S3d**); second, to ensure our ability to detect m^6^A variation, we silenced METTL14, an m^6^A writer. As anticipated, METTL14 targeting triggered a significant decrease of m^6^A level (but not m^6^A_m_) by LC-MS/MS (**Figure S4a**). Next, we performed a similar analysis with mRNA extracted from our panel of patients derived cell lines (**Figure 3b**). We observed an increased level of m^6^A_m_ / A ratio in CTC lines (**Figure 4b**) which was concomitant with low FTO levels (**Figure 3a**). By contrast, m^6^A / A ratio was rather decreased (**Figure 4b**). Importantly, FTO overexpression in two CTC lines triggered the opposite effect and decreased tremendously m^6^A_m_ / A ratio while m^6^A / A ratio remained unchanged (**Figure 4c**). Altogether, these observations established a first connecting thread between FTO-mediated m^6^A_m_ dynamic and the acquisition of cancer stem ability in colorectal cancer.

**Figure 4.**
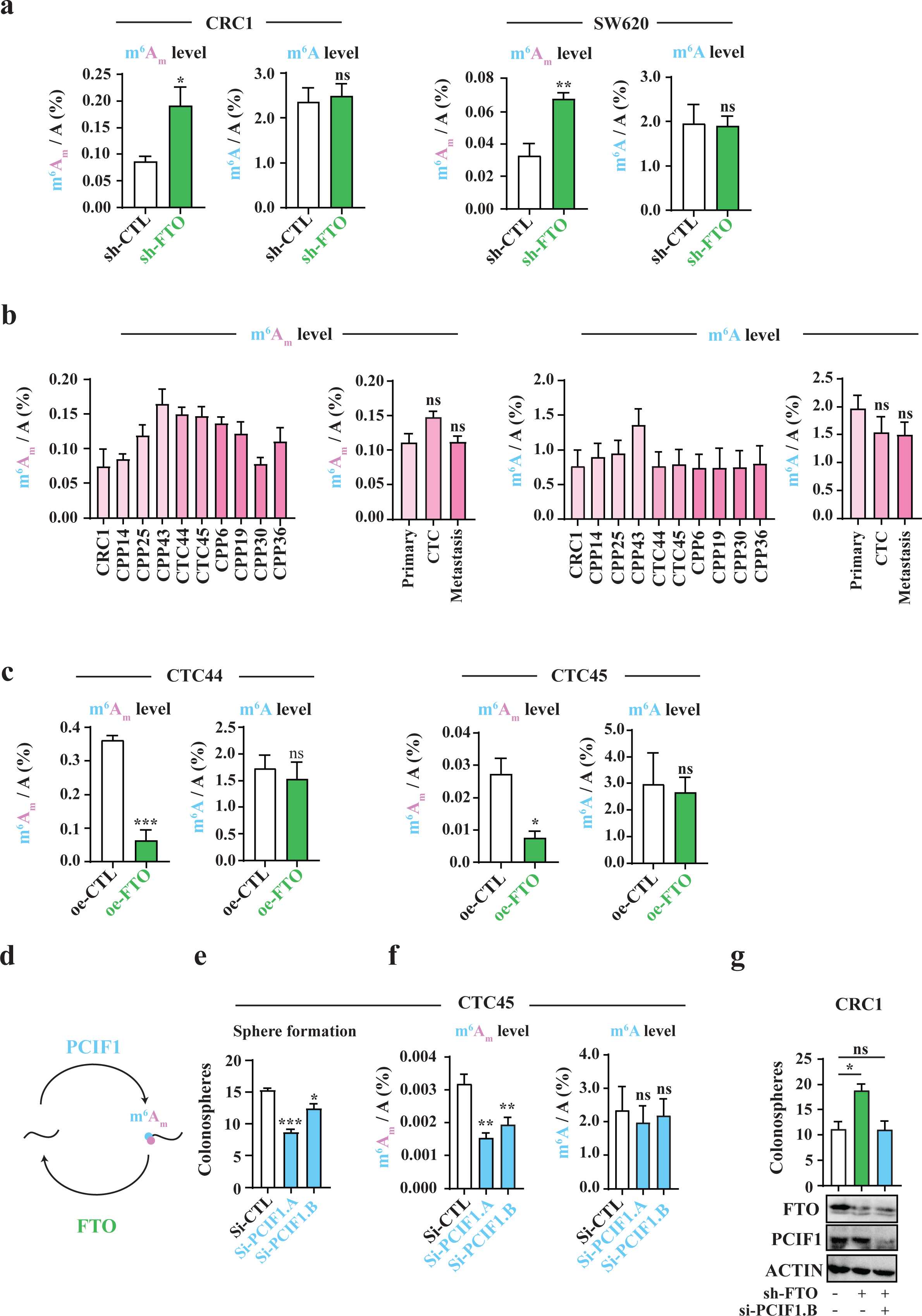
FTO affects stem-like properties through m^6^A_m_ demethylase activity. **(a) FTO silencing increases m**^**6**^**A**_**m**_ **level rather than m**^**6**^**A**. LC-MS/MS mRNA quantification of m^6^A_m_/A and m^6^A/A ratios in either CRC1 sh-FTO or sh-CTL and SW620 sh-FTO or sh-CTL cell line. Bar plot represents Mean +/- S.E.M of at least three biological replicates. ** p-value < 0.01, * p-value < 0.05, ns = not significant. Two-Sided Unpaired T-test. **(b) m**^**6**^**A**_**m**_ **tends to be increased in CTC lines but not m**^**6**^**A**. LC-MS/MS analysis of mRNA from patient derived cell lines (same as in **Figure 3**). m^6^Am/A and m^6^A/A were evaluated. Graphs represent mean +/- S.E.M of m^6^A_m_/A or m^6^A/A level for each cell line or group of cell lines. Three biological replicates. **(c) FTO overexpression decreases m**^**6**^**A**_**m**_ **rather than m**^**6**^**A in CTC lines**. mRNA quantification of m^6^A_m_/A and m^6^A/A after FTO overexpression in CTC44 line. Graphs represent mean +/- S.E.M of at least 3 biological replicates. *** p-value < 0.001, * p-value < 0.05, ns = not significant. Two-sided Unpaired T-test. **(d) Effectors of m**^**6**^**A**_**m**_ **modification. (e) PCIF1 silencing decreases colonosphere formation**. Sphere forming ability of CTC44 line after silencing of the m^6^A_m_ writer PCIF1. Bar plot represents mean +/- S.E.M of three experiments. *** p-value < 0.001, * p-value < 0.05. Two-sided Unpaired T-test. **(f) PCIF1 silencing decreases only m**^**6**^**A**_**m**_ **level**. LC-MS/MS mRNA quantification of m^6^A_m_/A and m^6^A/A after PCIF1 silencing in CTC44 line. Graphs represent mean +/- S.E.M of 3 biological replicates. *p-value < 0.05, ns = not significant. Two-Sided Unpaired T-test. **(g) PCIF1 silencing “rescues” sphere forming ability in CRC1 sh-FTO cell line**. PCIF1 was depleted by siRNA treatment in sh-FTO cell line and sphere formation assay was performed. Results are mean +/- S.E.M of three independent experiments. *p-value < 0.05, ns = not significant. One-way Anova followed by multiple comparisons.

Recent reports identified PCIF1/CAPAM as the m^6^A_m_ methyltransferase (“writer”) [28, 42] (**Figure 4d**) and proposed that PCIF1/CAPAM activity is involved in cellular resistance to oxidative stress response induction [28]. It is well known that elevated reactive oxygen species (ROS) production impairs self-renewal and promote cell differentiation in stem cells and their malignant counterpart [43]. To examine the contribution of PCIF1/CAPAM to CSC phenotype, we silenced PCIF1 in two CTC lines (**Figure S4b**) and tested both SFP and resistance to chemotherapy. As expected, PCIF1/CAPAM knockdown reduced sphere number by 30% (**Figure 4e, S4c**) as well as m^6^A_m_ / A ratio while m^6^A level remained unaffected (**Figure 4f, S4d**). Along the same vein, PCIF1 depletion in sh-FTO cells rescued the phenotype (**Figure 4g, S4e**). PCIF1 is also expressed at lower level in metastatic cell lines (**Figure S4f & g**), concomitantly with reduced FTO expression (**Figure 3a**). Balanced decrease of m^6^A_m_ writer and eraser (**Figure S4h**) would explain the lack of change of m^6^A_m_ level with respect to cell lines originating from primary tumor (**Figure 4b**).

Yet, PCIF1/CAPAM level did not affect CTC resistance to FIRI treatment (**Figure S5a**). Likewise, sub-lethal doses of FIRI, which promote CSC phenotype (**Figure 2d**), did not modify PCIF1 expression (**Figure S5b**). Nevertheless, these cells displayed reduced FTO expression (**Figure S5b**) as well as increased m^6^A_m_ level (**Figure S5c**). This suggests that FTO and PCIF1 do not exhibit mere antagonistic activities but rather share a partially overlapping enzyme substrate specificity. Distinct cellular distribution of these two effectors may explain such functional difference.

### FTO mediated m^6^A_m_ demethylation takes place in the cytoplasm

While FTO sequence carries a Nuclear Localization Signal (NLS), its cellular localization varies among several mammalian cell lines [39, 40]. This spatial regulation may result in distinct substrate preference: a recent report shows that FTO catalyzes both m^6^A and m^6^A_m_ demethylation of mRNA in cytoplasm but targets preferentially m^6^A in cell nucleus [41]. In our cell lines (CRC1 and SW620), FTO is present in nuclear speckles [17] as well as in cell cytoplasm (**Figure 5a**). We employed detergent-based cell fractionation procedure to separate nuclei from cytoplasm from sh-CTL and sh-FTO cells.

**Figure 5.**
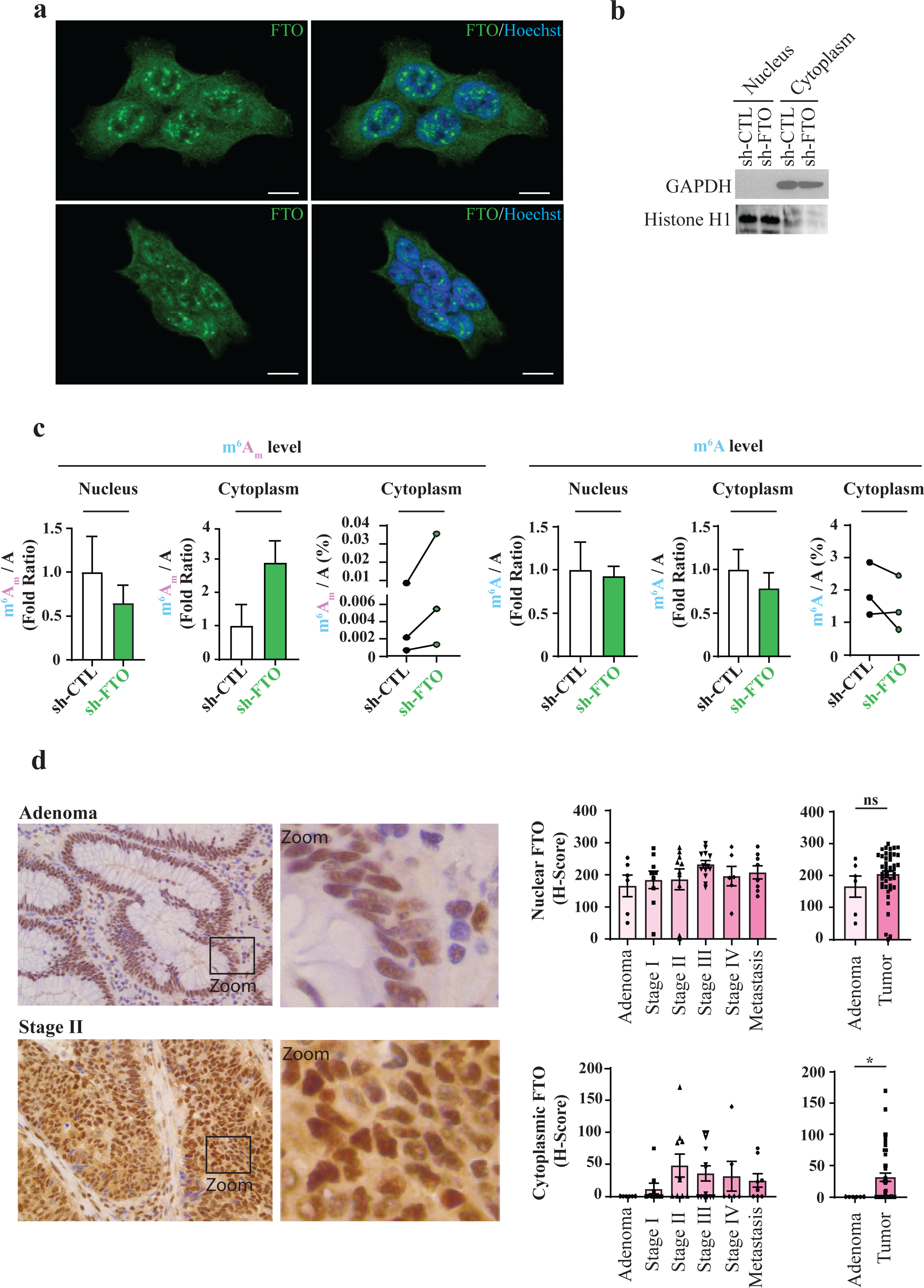
FTO mediated m^6^A_m_ demethylation takes place in the cytoplasm. **(a) FTO localizes both in nucleus and cytoplasm**. Immunofluorescence staining of FTO (green) in CRC1 and SW620 cell lines show presence of FTO in cytoplasm and in nuclear speckles. Nucleus of cells were stained with Hoechst (blue). Scale bar 10 μm. **(b) Verification of the effective cell fractionation procedure by immunoblot**. Effective separation of cytoplasmic and nuclear fractions was evaluated by immunoblot using cytoplasmic marker (GAPDH) and nuclear marker (Histone H1). **(c) FTO silencing increases cytoplasmic m**^**6**^**A**_**m**_ **level**. mRNA quantification of m^6^A_m_ / A and m^6^A / A level for nuclear fraction and cytoplasmic fraction. Bar plots represents mean +/- S.E.M of three biological replicates. Before – After plots represents the same data as bar plot with raw values. **(d) FTO relocalizes to the cytoplasm during tumorigenesis**. FTO expression and localization was evaluated by IHC on TMA from CRC patient. Then, nuclear and cytoplasmic level of FTO were quantified. Pictures are representative of stage 0 and stage 2. Bar plots represents mean +/- S.E.M of H-score based on nuclear and cytoplasmic intensity of FTO staining. Each dot corresponds to individual value. *p-value < 0.05, ns = not significant, Mann-Whitney test.

The efficacy of this protocol was evaluated by immunoblot using cytoplasm- and nucleus-specific markers, respectively GAPDH and Histone H1 (**Figure 5b**). Then, we extracted mRNA from both compartment and quantified m^6^A_m_ and m^6^A by LC-MS/MS (**Figure 5c**). We observed a significant increase (almost 3 fold) of m^6^A_m_ / A ratio in sh-FTO cytoplasm (in comparison with sh-CTL) while this ratio remained steady in the nucleus. By opposition, neither of these two cell compartments displays alteration of m^6^A / A ratio (**Figure 5c**). This suggests that FTO-mediated m^6^A_m_ demethylation takes place in the cytoplasm in colorectal cancer cell lines.

Next, we evaluated FTO expression and localization in tumor microarrays (TMA) from different colorectal stages: adenoma, 1, 2, 3, 4 and metastases (n=52). Global FTO expression did not show any significant change over the course of tumor evolution (**Figure 5d**). Yet, subcellular distribution analysis unveiled interesting features. FTO expression is strictly nuclear in healthy adjacent tissue as well as in precursor lesions of CRC (adenoma, **Figure 5d**). Then, following submucosal invasion (stage 1), FTO was also found in the cytoplasm (**Figure 5d**). These observations suggest that the tumorigenic process alters sub-cellular FTO distribution.

### FTO activity does not affect transcription process but rather translational control

FTO activity is associated with regulating post-transcriptional processes including mRNA splicing [44], stability [29] and translation [41]. To investigate the molecular mechanism of FTO regulation of colorectal CSCs we performed whole transcriptome sequencing (RNA-seq, 125 bp, paired-end, n=3) of sh-FTO and sh-CTL cells. The volcano plot of the transcriptomic variation between sh-CTL and sh-FTO cells (**Figure 6a**) reveals that only few mRNAs exhibit a fourfold change, and among those only 5 reach the standard log10(p –value) threshold of 1.5 (with n=3 replicates) for statistical significance: PUF60, GFOD1, PAK6, COL13A1 and BMP2K (**Table S4**).

**Figure 6.**
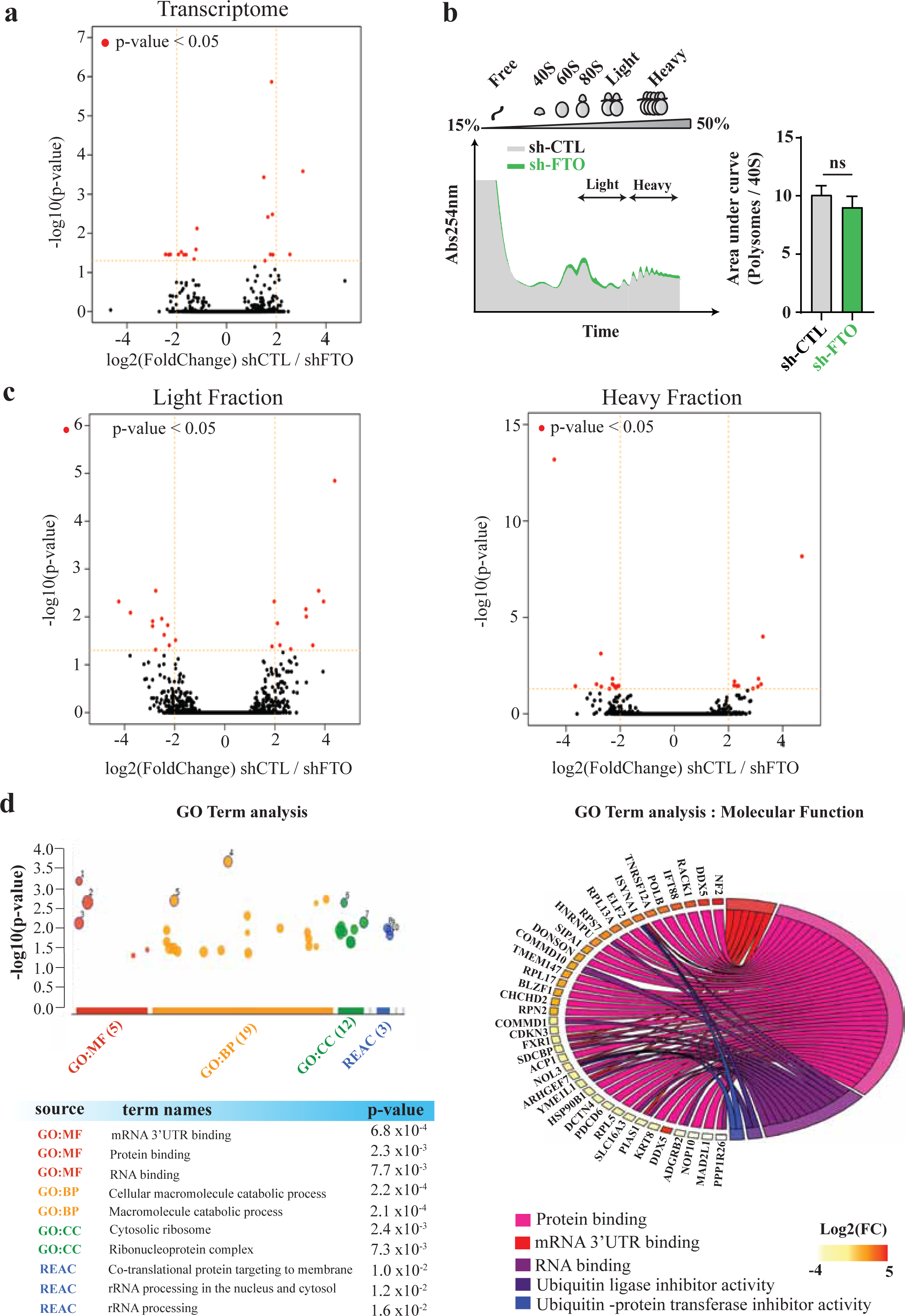
FTO silencing triggers subtle changes at the translational level. **(a) FTO silencing does not affect global transcript level.** MA plot analyses of transcripts changes. Red dot represent transcripts with an adjusted p-value < 0.05. Pie Chart shows the percentage of transcripts with fold change > 2 or < 0.5; CRC1 line n = 3 individual experiments. **(b) FTO does not affect global translation activity**. Polysome profile of sh-CTL and sh-FTO CRC1 cell lines. Light fraction (1 to 3 ribosomes/mRNA) and heavy fractions are indicated. Bar plot represents polysome level quantification of three experiments. ns = p-value > 0.05. **(c) FTO silencing impairs translation of specific mRNA**. MA plot analyses of transcripts changes in Light and Heavy fractions. Same representation as in **(a)**; CRC1 line, n = 3. **(d)** Gene ontology analysis. GO Term analysis was performed with gProfileR. Graph on the left shows significant GO terms of the three main categories (Molecular Function, MF; cellular compartment CC; Biological Process, BP). The table indicates the three most significant GO Terms for each category. MF GO Terms are detailed in the pie chart (right). Each color represents one GO Term that associates with the corresponding transcripts. Level of transcript changes is also indicated (below; right).

Such minor differences in mRNA levels suggest that FTO may act in CSCs by modulating translation efficiency of individual mRNAs. This scenario is consistent with the proximity of m^6^A_m_ adjacent to the m^7^G-Cap, as well as a recent study on PCIF1 suggesting a role of m^6^A_m_ modification in translational control of certain mRNAs [28]. Consistent with the lack of effect of FTO modulation on proliferation (**Figure S1e**), polysome fractionation profile did not reveal significant changes in global translation activity, which would be expected to modify the ratio between unassembled ribosome subunits, monosome and polysome size and numbers (**Figure 6b**).

We therefore performed RNA-seq on mRNA co-sedimenting with heavy polysomes (4+ ribosomes /mRNA) or light polysomes/monosomes (1-3 ribosomes per mRNA). We mapped reads to the human reference transcriptome to determine mRNAs that were differentially translated between sh-CTL and sh-FTO cell lines. The volcano plots of the light and heavy analyses (**Figure 6c**) show stronger changes in translation than in transcription (**Figure 6a**). Globally, 23 mRNAs were differentially translated in the heavy fraction, and 26 in the light fraction (|log2(FC)|>2 and (-log10(p-value)> 1.5), including ribosome related proteins (RPL13A, RPS7, FXR1, UTP4, or NOP10) (**Table S5 and S6**). We confirmed this selective translation engagement for several genes by qRT-PCR (**Figure S6**) Gene set enrichment analyses of differentially translated genes indicate their involvement in mRNA/macromolecule catabolic process (and regulation), in apoptotic process, and molecular functions such as mRNA 3’UTR binding (**Figure 6d**). Analysis with gProfiler indicated three altered pathways, all related to translation: rRNA processing in the nucleus and cytosol and SRP-dependent co-translational protein targeting to membrane (Reactome ids: R-HSA-72312, R-HSA-8868773, R-HSA-1799339). Taken together, these results point to an impact of FTO activity on the fine-tuning of translation process in colorectal cancer cells.

### m^6^A_m_-modified transcripts are enriched in translationally up-regulated genes

Next, we examined whether the FTO effect on translational control could be mediated by m^6^A_m_ modification. We used data from Wei *et al*. [41] to identify m^6^A_m_ modified transcripts among either transcriptionally or translationally regulated genes. We could not see any enrichment of m^6^A_m_ genes in the shortlist of transcriptionally regulated mRNAs (**Figure 7a**). However, m^6^A_m_ marked genes were significantly enriched in the regulated mRNA populations of mRNA from light and heavy fractions (**Figure 7b**). The cumulative distribution of transcripts correlated m^6^A_m_, but not m^6^A marking, with enhanced translation (**Figure 7c**) as previously suggested by Akichika *et al*. [28]. Accordingly, FTO silencing triggers enrichment of m^6^A_m_-modified transcripts in upregulated genes but not in downregulated ones (**Figure 7d**).

**Figure 7.**
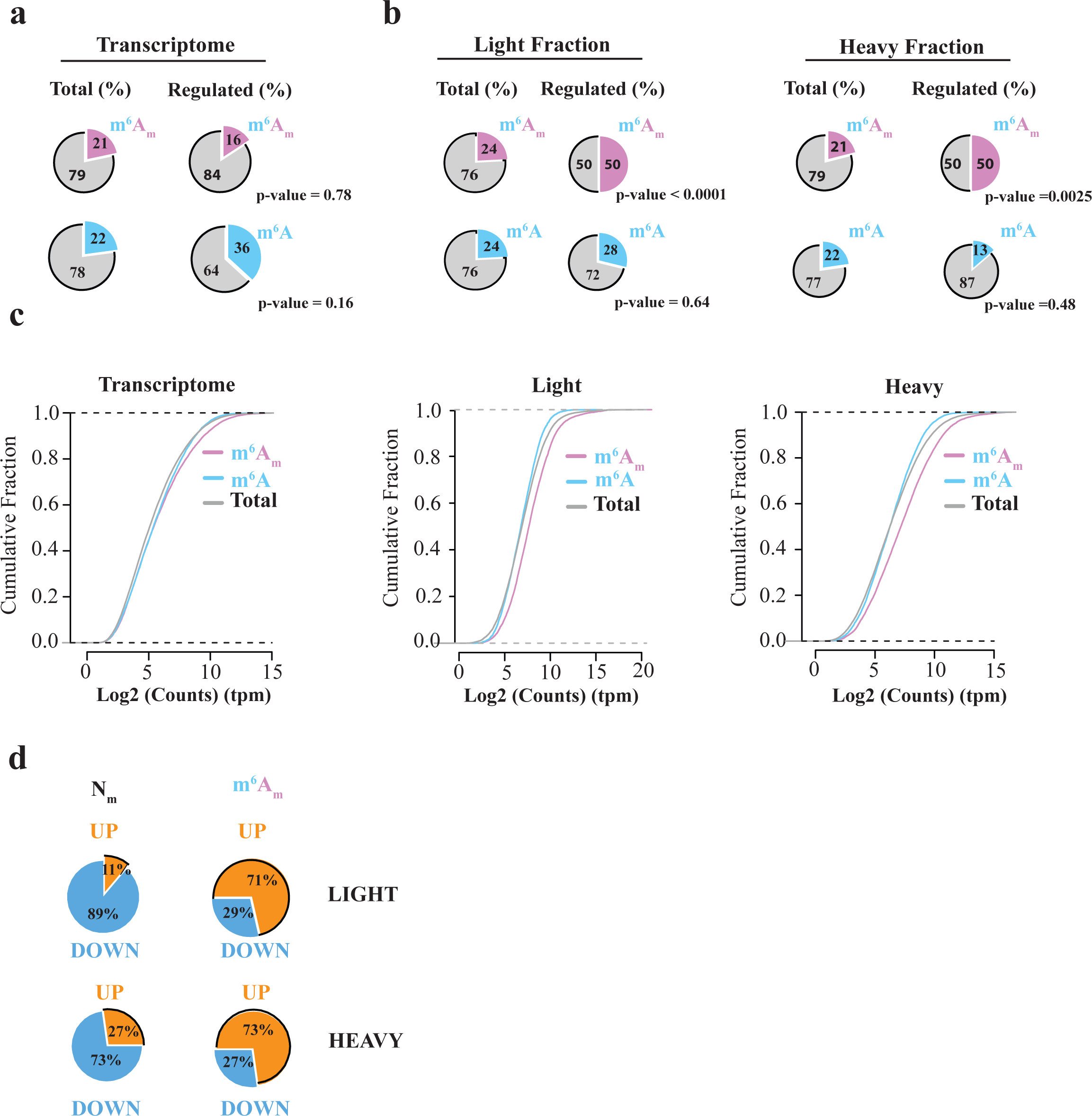
m^6^A_m_ marking correlates with enhanced translation. **(a) m**^**6**^**A**_**m**_ **marking does not correlate with transcription regulation**. Fraction of potential m^6^A_m_ marked transcripts (identified by Wei J *et al. [41]*) in total mRNA and significantly regulated transcripts (sh-FTO CRC1 cell line, FDR < 0.05). P-values were obtained using Fisher exact test. **(b) Translationally regulated transcripts are enriched in m**^**6**^**A**_**m**_ **transcripts**. Same as in **(a)** with translatome data from light and heavy fractions (sh-FTO CRC1 cell line, FDR < 0.05). P-values were obtained using Fisher exact test. **(c) m**^**6**^**A**_**m**_ **transcripts are enriched in polysome fractions from CRC1 cell line**. Cumulative distribution plot that represents the fold change of mRNA that starts with either m^6^A_m_ or m^6^A from sh-FTO vs sh-CTL cells. Both m^6^A_m_ (pink) and m^6^A (blue) transcripts were identified based on data from Wei J *et al*. The level of m^6^A_m_ transcripts is more important in polysome fractions (light and heavy) in comparison with total transcripts. Graph represents the number of counts for m^6^A_m_, m^6^A and total transcripts in transcriptomic data, light and heavy fractions. n = 3, CRC1 cell line. **(d) m**^**6**^**A**_**m**_ **marking tends to favor mRNA translation**. Translationally deregulated transcripts were divided into two categories: upregulated or downregulated genes. Most of m^6^A_m_ modified transcripts belong to the upregulated category. Other transcripts (Nm) show the opposite effect.

## DISCUSSION

Our study is the first to address the specific function of FTO in recently established colorectal cancer cell lines with a focus on CSC phenotype. We show that decreased FTO activity plays a critical role in colorectal cancer by enhancing CSC properties including sphere forming, *in vivo* tumorigenesis and chemoresistance. The underlying mechanism takes place in the cytoplasm and appears to involve demethylating m^6^A_m_ residues adjacent to the m^7^G-cap in highly selected transcripts, most likely to adjust their translation efficiency.

As the initially characterized m^6^A demethylase, FTO has been studied in various types of cancers, often reported as a pro-oncogenic factor [45]. Inhibiting FTO in glioblastoma impairs self-renewal and cancer progression [46]. FTO is highly expressed in some AML types where it suppresses all-trans retinoic acid-induced cell differentiation and promotes oncogene-mediated cell transformation and leukemogenesis [45]. High levels of FTO are characteristic in cervical squamous cell carcinomas where it promotes resistance to chemo/radiotherapy and increases DNA damage responses [47]. More recently, high FTO expression was associated with lower survival rates in patients with breast cancer [48]. Finally, FTO promotes breast cancer cell proliferation, colony formation and metastasis *in vitro* and *in vivo* [48]. By contrast, our data support an anti-oncogenic role of FTO in colorectal cancer, emphasizing the importance of tissular context. This observation is consistent with previous studies showing cancer-specific effects of m^6^A regulators. METLL3 activity promotes tumorigenesis in AML by enhancing BCL2 and PTEN translation [49], whereas in glioblastoma stem cells, it suppresses growth and self-renewal by reducing expression of ADAM19 [50].

Interestingly, Kaplan-Meier survival analysis from cancer database shows that overall survival of colorectal cancer is higher with lower FTO. This is contradictory with our data that connects reduced FTO expression with enhanced chemoresistance and tumor initiation, as emphasized by our results from cell lines derived from CTCs, the source of lethal metastases. At least two parameters can account this inconsistency. First, we show that FTO expression in colon cancer cells is finely regulated at the post-transcriptional level, though the precise underlying mechanism remains to be determined. This observation is in agreement with a recent report in gastric tissue [38] and stresses the importance of quantifying gene expression at the protein level for diagnostic purpose. Second, as will be discussed below, FTO activity and substrate specificity may vary with its subcellular distribution. This parameter is even more critical in the context of colorectal cancer where a fraction of FTO relocate from the nucleus to the cytoplasm at early stage of the disease. Whether the cause may be due to neoplastic transformation process remains to be determined.

Despite being ubiquitously expressed, FTO activity and function vary widely among tissues, as illustrated by transcriptome-wide mapping of m^6^A modifications [25, 26]. To better comprehend the biological function of FTO, extensive efforts have been made to identify relevant RNA substrate(s). No clear consensus emerges from recently published studies [29, 41, 51, 52]. Following the identification of FTO as the first m^6^A mRNA demethylase [17], several reports connected demethylation of internal m^6^A nucleotides with a wide range of biological processes such as viral infection, stress- and DNA UV damage-responses [53-55]. More recently, Mauer *et al*. threw a stone into the pond by suggesting that FTO preferentially demethylates m^6^A_m_ residues closely adjacent to the cap [29]. The higher *in vitro* demethylation efficiency reported was confirmed by Wei *et al*., although the conclusion was moderated in the cellular context, since the level of internal m^6^A marking is ten-fold higher than m^6^A_m_ marking [41]. Discrepancies between studies on FTO may arise from several known biological parameters. A recent review addresses this ambiguity and emphasize the context-dependent functions of RNA methylation effectors [56]. First, FTO displays a wide substrate spectrum: besides mRNA (and tRNA for m^1^A [41]), FTO can demethylate m^6^A and m^6^A_m_ in snRNAs [51]. Second, its spatial distribution dictates substrate preferences: cytoplasmic FTO catalyzes demethylation of both m^6^A and cap-m^6^A_m_ whereas nuclear FTO demethylates preferentially m^6^A [41], most likely owing to accessibility constraints to the cap-moiety. In colorectal cancer cell lines, FTO activity displays a remarkable selectivity for cap-m^6^A_m_ which concurs with its presence in the cytoplasm. LC-MS/MS analysis of small RNA species did not reveal any change of m^6^A and m^6^A_m_ levels following FTO silencing. Also, our bioinformatic analysis of sh-FTO vs sh-CTL RNA-seq data failed to identify significant alterations of mRNA splicing that might result from altered snRNA methylation. While we cannot exclude subtle alteration of internal m^6^A marks, our data clearly establishes cap-m^6^A_m_ residues of mRNA as the main substrate of cytoplasmic FTO in colorectal cancer cells. Extending this observation, targeting either the m^6^A writer complex or ALKBH5 does not significantly affect CSC phenotype. By contrast, inhibiting PCIF1/CAPAM partially recapitulated the inhibiting effect of FTO overexpression on the CSC phenotypes. PCIF1/CAPAM catalyzes cap-A_m_ methylation in the nucleus while FTO demethylates m^6^A_m_ in the cytoplasm. Compartment-specific enzymatic activity implies partially overlapping mRNA substrates and explains why m^6^A_m_ writer and eraser do not display mere antagonistic activities.

In this study, we clearly identify m^6^A_m_ as a critical epitranscriptomic mark for controlling the stem cell phenotype of human colon cancer cells. While m^6^A_m_ was first identified in mRNA from mammalian cells and viruses in the 70s [27], the enzymes catalyzing m^6^A_m_ modifications, PCIF1/CAPAM and FTO, were only recently identified, and the study of m^6^A_m_ role in mRNA metabolism and cellular function is still in its infancy. Due to its abundance and its strategical location in the cap structure [57], this chemical modification holds an inherent potential in gene expression control. Mauer *et al*. reported that FTO-catalyzed m^6^A_m_ demethylation reduced mRNA stability by rendering it more vulnerable to DCP2-mediated decapping process [29]. While our study supports the function of FTO as m^6^A_m_ demethylase, we noticed minor changes at the transcriptome level following FTO knockdown. Yet, FTO depletion enhanced translation of known m^6^A_m_ marked transcripts, which concurs with the study from Akichika *et al*. that connected loss of m^6^A_m_ modification in PCIF1/CAPAM knockout cells with decreased translation of mRNA [28]. Several deregulated genes relate to translation machinery (e.g. RPL13A, RPS7, FXR1, UTP4, or NOP10) which underscores the role of FTO/m^6^A_m_ axis in fine-tuning the translation process. Whether this involves subtle change in the composition of the translation machinery, that would alter its ability to filter mRNA [58], remains to be assessed. Nevertheless, none of the key regulatory genes generally associated with CSC traits, such as the octamer-binding transcription factor 4 (OCT4), SOX2 and NANOG [59, 60], were altered in sh-FTO cells. This suggests that, in colorectal cancer cell, FTO-mediated m^6^A_m_ demethylation restrains CSC abilities through reversible, unconventional mechanism that may involve reprogramming of gene expression. Nevertheless, while we see modest but significant differences at the translation level, we cannot exclude the involvement of other mechanism(s).

In summary, we have shown that cytoplasmic FTO activity regulates m^6^A_m_ modification of selected mRNAs and subtly but surely is necessary for maintaining the CSC phenotype *in vitro* and *in vivo* for human colon cancers. The extent to which this applies to other cancers remains to be determined: given the complexity of cancer, there will likely be a great variability between cancer types. Even if limited to colon cancer, however, our findings point the way to deeper understanding of the CSC phenotype and have exciting implications for developing therapies that decrease m^6^A_m_ modification to cripple CSC-based metastases and resistance.

## ONLINE CONTENT

The online version of the paper contains additional methods, along with supplementary figures and tables that support the findings. References unique to these sections appear only in the online paper.

**SUPPLEMENTARY INFORMATION** is available in the online version of the paper.

## ACKNOWLEDGEMENTS

This work was generously supported by Ligue contre le Cancer, SIRIC Montpellier Cancer *(INCa-DGOS-Inserm 6045)*, Labex NumeV (GEM flagship project) and Occitanie Region/FEDER (PPRi, SMART project). We thank Montpellier Genomix (http://www.mgx.cnrs.fr) sequencing facility as well as iExplore animal facility, in particular Denis Greuet and Steeve Thirard. We thank the ATGC bioinformatic platform, whose servers hosted our bioinformatic analyses, and which belongs to the “France Génomique” network and to Institut Français de Bioinformatique.

The authors declare no conflict of interest.

## AUTHOR CONTRIBUTIONS

A.D., A.B., E.R., S.R. designed experiments and analysed the results. S.R., H.G., A.Am., F.B., A.At., J.V., F.D., A.C., F.M. performed experiments. A.D., A.B., E.R., S.R., C.H., E.C., JJ.V., E.S., J.P. performed data analyses. J.R. and S.R. designed bioinformatics pipelines and performed bioinformatics analysis. A.D., A.B., E.R., S.R. wrote the manuscript. All the authors reviewed the final version of the manuscript.

## METHODS

### Cell lines

Patient-derived colon cancer cell lines (CRC1, CPP-14/25/43/6/19/30/36) were derived from CRC surgeries provided by CHU-Carémeau (Nîmes, France, ClinicalTrial.gov Identifier#NCT01577511) within an approved protocol. CRC1, CPP-14/25/43 cell lines were derived from primary tumors and CPP-6/19/30/36 from metastatic tumors. CTC44 and CTC45 are circulating tumor cell lines derived from blood of metastatic chemotherapy-naïve stage IV CRC patients [31]. HCT-116 (ATCC^®^ CCL-247™) and SW620 (ATCC^®^ CCL-227™) are commercially available colon cancer cell lines derived respectively from primary and metastatic human tumors.

### Cell culture and generation of stable cell lines

Cells were maintained at 37 °C under humidified 5% CO_2_ in DMEM medium supplemented with 10% FCS (Invitrogen) and 2 mM glutamine. Stable knockdown of FTO was achieved by lentiviral delivery (5 D.O.I) of anti-FTO sh-RNA (Origene, #TL308064, sh-FTO#B). Isolation of infected cells was performed by GFP positive cells sorting on FACSAria.

### Plasmid constructions

FTO coding sequence was amplified from pDONOR plasmid (Montpellier Genomic Collection MGC Facility) and inserted into pCDNA3-Flag Cter plasmid into *Hind*III and *Xba*I restriction sites.

### Transfections

Transfection of 100 nM of si-RNA duplex was performed using Lipofectamine RNAimax (Invitrogen) according to the manufacturer’s instructions. The sequences of siRNA are provided in **Table S3**. 300,000 cells in 6-well plate were transfected for 48 h with 2 µg of plasmid DNA using Lipofectamine 2 000 (Invitrogen) according to the manufacturer’s instructions.

### FIRI treatment

Cells were treated for 72 h at 37 °C under humidified 5% CO2 with 10 µM of 5-Fluorouracile coupled with 0.1 µM of SN38.

### RNA extraction and RT-qPCR

Total RNA was extracted using TRIzol reagent (Invitrogen) according to the manufacturer instructions. For RT-qPCR analyses, 1 µg of RNA was reverse-transcribed into cDNA using random hexamer (Invitrogen) and 1 U of MML-V reverse transcriptase (Invitrogen). Quantitative gene expression was performed using SYBR Green master mix (Roche) on LightCycler 480 Instrument (Roche). Results were normalized to actin expression and analyzed using the ΔΔCt method. Primer sequences are provided in **Table S1**.

### Protein extraction and western blot

Cells were washed twice with ice cold phosphate-buffered saline (PBS) and lysed in RIPA buffer (50 mM Tris, 150 mM NaCl, 1% NP-40, 0.25% sodium deoxycholate, 2 mM sodium orthovanadate, 50 mM Sodium fluoride, 50 mM β-glycerophosphate, 2 mM EGTA; pH 7.5). Samples were separated on a 12% SDS–polyacrylamide gel electrophoresis, transferred to nitrocellulose membrane, blocked for 1 h in 5% (w/v) non-fat dry milk in PBS and probed overnight at 4 °C with primary antibodies (**Table S2**). Membranes were incubated with secondary antibody for 1 h and proteins were revealed by ECL Prime (Amersham) using ChemiDoc Touch imager (Biorad).

### Proliferation assay

1,000 cells were seeded in triplicate in 96-well plate for 24, 48, 72 and 96 h. Cells were fixed for at least 2 h at 4 °C in 10% TCA. After three washes in miliQ water, cells were incubated in 0.4% Sulforhodamine B solution for 30 min at room temperature followed by three washes with miliQ water. 562nm absorbance was measured after resuspension in 10 mM Tris-Base.

### Sphere formation assay

This test was performed as previously described [61]. Number of sphere forming cells were determined after plating of 100 cells / 100 µL of M11 medium (DMEM/F12 (1:1) Glutamax medium, N2 Supplement, Glucose 0.3 %, insulin 20 µg/ml, hBasic-FGF 10 ng/ml, hEGF 20 ng/ml) in ultra-low attachment 96 well-plates. Spheres > 50 µm were counted after 5 – 7 days.

### Cytotoxicity assay

2,000 cells were seeded in a 96-well plate. After 24 h, cells were treated with decreasing doses of 5-Fluorouracile (5-Fu) coupled with SN38 (FIRI) for 72 h (1/3 dilution from 3.3 X to 0 X; 1 X = 50 µM 5-FU + 0.5 µM SN38). FOX toxicity was assessed with decreasing doses of 5-Fluorouracile coupled with oxaliplatin for 72 h (1/3 dilution from 3.3 X to 0 X; 1 X = 50 µM 5-FU + 1µM oxaliplatin). Cell viability was measured using Sulforhodamine B assay as previously described and IC_50_ was determined graphically.

### Flow cytometry

The ALDH activity of adherent cells was measured using the ALDEFLUOR kit (Stem Cell Technologies), according to the manufacturer’s instructions. CD44 and CD44v6 were stained using anti-CD44 antibody and anti-CD44v6 antibody for 15 min at 4 °C (**Table S2**). As a reference control, anti-IgG2a and REA-S control Isotype was used under identical conditions. The brightly fluorescent ALDH, CD44 or CD44v6 positive cells were detected using a MACSQuant Analyzer (Miltenyi Biotec). To exclude nonviable cells, Sytox blue was added at a concentration of 0.1 µg/ml.

### Immunofluorescence

Cells were fixed in PBS containing 4% paraformaldehyde at room temperature for 15 min, wash twice in PBS, permeablized with 0.1% NP-40 in PBS for 10 min, wash twice in PBS and blocked with 5% FCS for 30 min. Coverslips were incubated 1h with primary antibody (Phosphosolution, 597-FTO) at RT. After washing three times with PBS, coverslips were incubated for 1 h with Alexa Fluor^®^-conjugated secondary antibody (Alexa Fluor^®^ 488 Goat Anti-Mouse (IgG), Invitrogen) at RT. For nuclei staining, coverslips were washed twice and incubated with 1 µg/ml Hoechst 3358 for 5 min at RT. After two washes with distilled water, coverslips were mounted on slides with Fluoromount-G (Invitrogen). Fluorescent pictures were acquired at room temperature on an AxioImager Z1 microscope (Carl Zeiss, Inc.) equipped with a camera (AxioCam MRm; Carl Zeiss, Inc.) and Plan Apochromat (63×, NA 1.4) objective, the Apotome Slider system equipped with an H1 transmission grid (Carl Zeiss, Inc.), and Zen 2 imaging software (Carl Zeiss, Inc.).

### mRNA purification

mRNA was purified from total RNA with 2 rounds of GeneElute mRNA purification kit (Sigma). rRNA was removed using Ribominus kit (Invitrogen) according to the manufacturer’s instructions.

### Nucleoside mass-spectrometry analysis

This part was performed as previously described [41]. Briefly, 400 ng of RNA was digested by 5 U of RppH (New England Biolabs) for 2 h at 37 °C. Decapped mRNA were then digested by 1 U of Nuclease P1 (Sigma) for 2 h at 42 °C in NH_4_OAc buffer (10 mM, pH 5.3). Nucleotides were dephosphorylated for 2 h at 37 °C by 1 U of Alkaline phosphatase in 100 mM of NH_4_OAc. The sample was then filtered (0.22 μm pore size, 4 mm diameter, Millipore), and 10 μL of the solution was injected into LC-MS/MS. The nucleosides were separated by reverse phase ultra-performance liquid chromatography on a C18 column with online mass spectrometry detection using Agilent 6490 triple-quadrupole LC mass spectrometer in multiple reactions monitoring (MRM) positive electrospray ionization (ESI) mode. The nucleosides were quantified by using the nucleoside-to-base ion mass transitions of 282.1 to 150.1 (m^6^A), 268 to 136 (A), 296 to 150 (m^6^A_m_) and 282 to 136 (A_m_).

### Tumor initiation assay

Decreasing amount of cells (1,000, 500, 100) were subcutaneously injected into nude mice (Hsd:Athymic Nude-Foxn1nu nu/nu, 6 weeks, females, 5 mice per group) in Matrigel - DMEM (v : v). Tumor sizes were measured twice a week for 50 days. After 50 days, the mice were sacrificed and tumors were taken out. The number of mice bearing growing tumor (size > 100 mm^3^) was counted. Tumor apparition frequency was determined using online ELDA (extreme limiting dilution analysis) software (https://bioinf.wehi.edu.au/elda/software).

### Chemoresistance *in vivo*

50,000 cells were subcutaneously injected into nude mice in Matrigel – DMEM (v : v). 50mg/kg 5-FU + 30mg/kg Irinotecan treatment (two i.p injection a week), was initiated once tumor reached 100 mm^3^ [62]. These studies were approved by the ethics committee of the Languedoc Roussillon Region and carried out in compliance with the CNRS and INSERM ethical guidelines of animal experimentation (CEEA-LR-12051).

### Purification of nucleus and cytoplasm

Cells were washed twice in ice-cold PBS. Cells were resuspended in fractionation buffer (0.5% NP-40, 150 mM NaCl, 50 mM Tris-HCl pH 7.4) and left on ice for 3 min. Cytoplasms were then extracted and nuclei were washed twice in wash buffer (150 mM NaCl, 50 mM Tris-HCl pH 7.4). Nuclei were resuspended in fractionation buffer and left on ice for 2 h. Cell debris were removed by centrifugation at 13 200 rpm for 10 min at 4 °C.

### Tissue Microarray (TMA)

Tissue microarray was constructed with FFPE tumor samples collected in the frame of the Clinical and Biological Database BCBCOLON (registered at ClinicalTrials.gov as NCT03976960). Adenomas, primary adenocarcinomas and metastatic lesions were sampled as two cores of 1mm diameter. All samples were chemo-naive. Tumor samples were collected following French laws under the supervision of an investigator and declared to the French Ministry of Higher Education and Research (declaration number DC-2008–695). The study was approved by the local translational research committee (ICM-CORT-2018-28).

### FTO detection by immunohistochemistry

Three-µm thin sections of formalin-fixed paraffin-embedded tissues were mounted on Flex microscope slides (Dako) and allowed to dry overnight at room temperature before immunohistochemistry processing, as previously described [63].Briefly, PT-Link^®^ system (Dako) was used for pre-treatment. Then, heat-induced antigen retrieval was executed for 15 minutes in High pH Buffer (Dako) at 95°C. Immunohistochemistry procedure was performed using the Dako Autostainer Link48 platform. Endogeneous peroxidase was quenched using Flex Peroxidase Block (Dako) for 5 min at room temperature. Slides were then incubated with the anti-FTO rabbit monoclonal antibody (AbCam, Clone EPR6895; 1/1000) for 20 min at room temperature. After two rinses in buffer, the slides were incubated with a horseradish peroxidase-labeled polymer coupled to secondary anti-mouse and anti-rabbit antibodies for 20 min, followed by appliance of 3,3’-Diaminobenzidine for 10 min as substrate. Counterstaining was performed using Flex Hematoxylin (Dako) followed by washing the slides under tap water for 5 min. Finally, slides were mounted with a coverslip after dehydratation.

Two independent observers analyzed the TMA slide in a blinded manner. The semi quantitative H-score method [64] was used to convert the expression of FTO to continuous values, based on both the staining intensity and the percentage of cells at that intensity. Nuclear and cytoplasmic signals were taken into account separately. Staining intensity was scored as no staining (0), weak staining (1), moderate staining (2), or intense staining (3). The percentage of cells stained at certain intensity was determined and multiplied by the intensity score to generate an intensity percentage score. The final staining score of each tissue sample was the sum of the four intensity percentage scores, and these scores ranged from 0 (no staining) to 300 (100% of cells with intense staining). In all cases with discrepant results, a consensus was reached between both investigators. Averaged FTO H-score was given when both cores from a single sample were assessable.

### Transcriptome Analysis

Total RNA was extracted using TRIzol method. RNA samples were send to Eurofins Genomics /GATC to perform next generation sequencing on Illumina platform.

### Polysome fractionation

This procedure was performed as previously described [65]. Briefly, 6 plates (150 cm^2^) were seeded with 2 × 10^6^ cells. After 72 h, cells were treated with 20 µg / ml emetine for 5 min at 37°C, washed twice with ice-cold PBS, and scraped in ice cold PBS. Cells were centrifuged, resuspended into 1mL of polysome lysis buffer, homogenized by hard shaking with 1,4mm ceramix spheres (Lysing matrix D MPBio) in FastPrep machine (MPBio), centrifuges 10 min at full speed at 4°C. Lysates were loaded on 15-50% sucrose gradient and centrifuged at 35,000 rpm for 2.5 h at 4°C in a SW41 rotor (Beckman Coulter). Polysomes were separated through a live optical density (OD) 254 nm UV spectrometer and collected with an ISCO (Lincoln, NE) density gradient fractionation system. The absorbance at 254 nm was measured continuously as a function of gradient depth.

### Translatome Analysis

mRNA associated with 1 to 3 ribosomes “Light polysomes” and mRNA associated with more than 3 ribosomes “Heavy polysomes” were extracted using TRIzol LS (Invitrogen) according to manufacturer’s instructions. Library preparation was performed using TruSeq Stranded mRNA Sample Preparation kit (Illumina) according to the manufacturer’s protocol. cDNA libraries were sequenced using the sequencer HiSeq 2500 (Illumina).

### Bioinformatic pipeline

Transcriptome and translatome libraries read quality were assessed using FastQC v0.11.5 (Babraham Institute, Cambridge, UK). Ribosomal RNA were discarded using SortMeRNA v2.1b **[66]**. High quality reads were then aligned on the *Homo sapiens* reference transcriptome, version GRCh38.cdna, and quantified using pseudocounts with Kallisto v0.45.0 **[67]**. Kallisto quantification parameters were fixed at 25 for k-mer size for the index, 20 for standard deviation and 100 for bootstraps. Statistical differential analyses were performed on each dataset using Wald test from DESeq2 R package **[68]**. Each count dataset was filtered at 1 count per millions per biological sample after size factors estimation, then dispersion was estimated. Primary risk of probabilities to false discovery fold change was corrected by Benjamini and Hochberg multiple test adjustment. Corrected p-values < at 0.05 % were kept. Volcano plots were realized with ggplot2 R package **[69]**. Gene identifications were performed with biomaRt R package **[70]**. Functional annotations were performed with online gProfileR **[71]** using a g:SCS threshold < at 0.05.

## DATA AVAILABILITY

Data collection will be freely available by DOI hosted in DRYAD (https://datadryad.org/stash) and/or on NCBI BioProject. Three links will be provided soon, one for transcriptome data, one for light translatome fraction and one for heavy translatome fraction.

## CODE AVAILABILITY

Data analysis was performed using free software detailled in the material and methods (i.e. FastQC v0.11.5, SortMeRNA v2.1b, Kallisto v0.45.0 and R v3.5.1). Statistics and graphics were performed using R packages (DESeq2, ggplot2, biomaRt) and an online program gProfileR. All the scripts used are hosted on a private gitlab depository and could be available from the corresponding author on reasonable request (contact).

## BIOLOGICAL MATERIAL AVAILABILITY

Any material used in this study is available from commercial source (indicated in the Materiel and Method section) or from the authors upon request.

