## Supplemental data for "FTO-mediated cytoplasmic m^6^A_m_ demethylation adjusts stem-like properties in colorectal cancer cell"

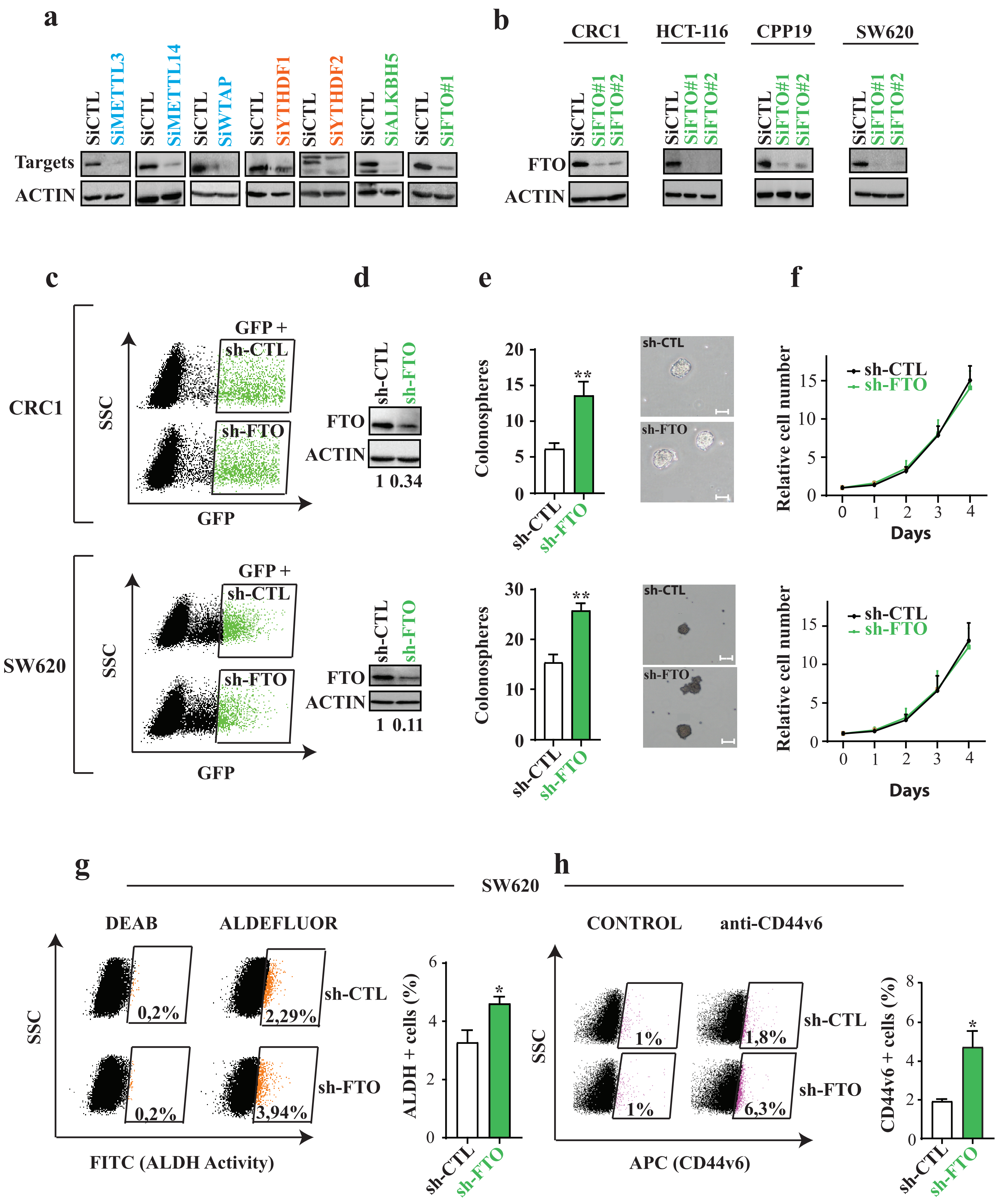

a
SiCTL
SiMETTL3
SiMETTL14
SiWTAP
SiYTHDF1
SiYTHDF2
SiALKBH5
SiFTO#1
Targets
ACTIN
b
CRC1
HCT-116
CPP19
SW620
SiFTO#1
SiFTO#2
FTO
c
CRC1
SW620
GFP +
sh-CTL
sh-FTO
SSC
GFP
d
1 0.34
1 0.11
e
Colonospheres
f
Relative cell number
Days
g
SW620
DEAB
ALDEFLUOR
0,2%
2,29%
3,94%
FITC (ALDH Activity)
ALDH + cells (%)
h
CONTROL
anti-CD44v6
1%
1,8%
6,3%
APC (CD44v6)
CD44v6 + cells (%)

**Figure S1. (a-b) Depletion of $m^6A$ actors by siRNA. (a)** Western blot analysis was performed to check the depletion of the $m^6A$ -related proteins. **(b)** Control of FTO depletion in 4 cell lines. Pictures are representative of three experiments. **(c - d) Establishment of stable cell lines. (c)** Efficiently infected cells by sh-RNA retrovirus were sorted by FACS on the basis of GFP expression (CRC1 top, SW620 bottom). **(d)** Efficient FTO depletion was evaluated by immunoblot. **(e) Stable FTO silencing increases sphere forming potential of CRC cell lines.** Colonosphere number after FTO silencing in CRC1 (top) and SW620 (bottom) cell lines. Results represent mean +/- S.E.M of three experiments. *** p-value < 0.001 ; ** p-value < 0.01, Two-sided Unpaired T-test. **(f) FTO silencing does not affect cell proliferation.** Cell proliferation was measured every day for 4 days using sulforhodamine assay (CRC1 top, SW620 bottom). Results are mean +/- S.E.M of three experiments. ns = not significant, two way ANOVA test. **(g–h) Stable FTO silencing increases CSC markers in SW620 cell line.** Number of ALDH (left) and CD44v6 (right) positive cells were assessed by Flow cytometry. Results are mean +/- S.E.M of three experiments. * p-value < 0.05, Two-sided Unpaired T-test.

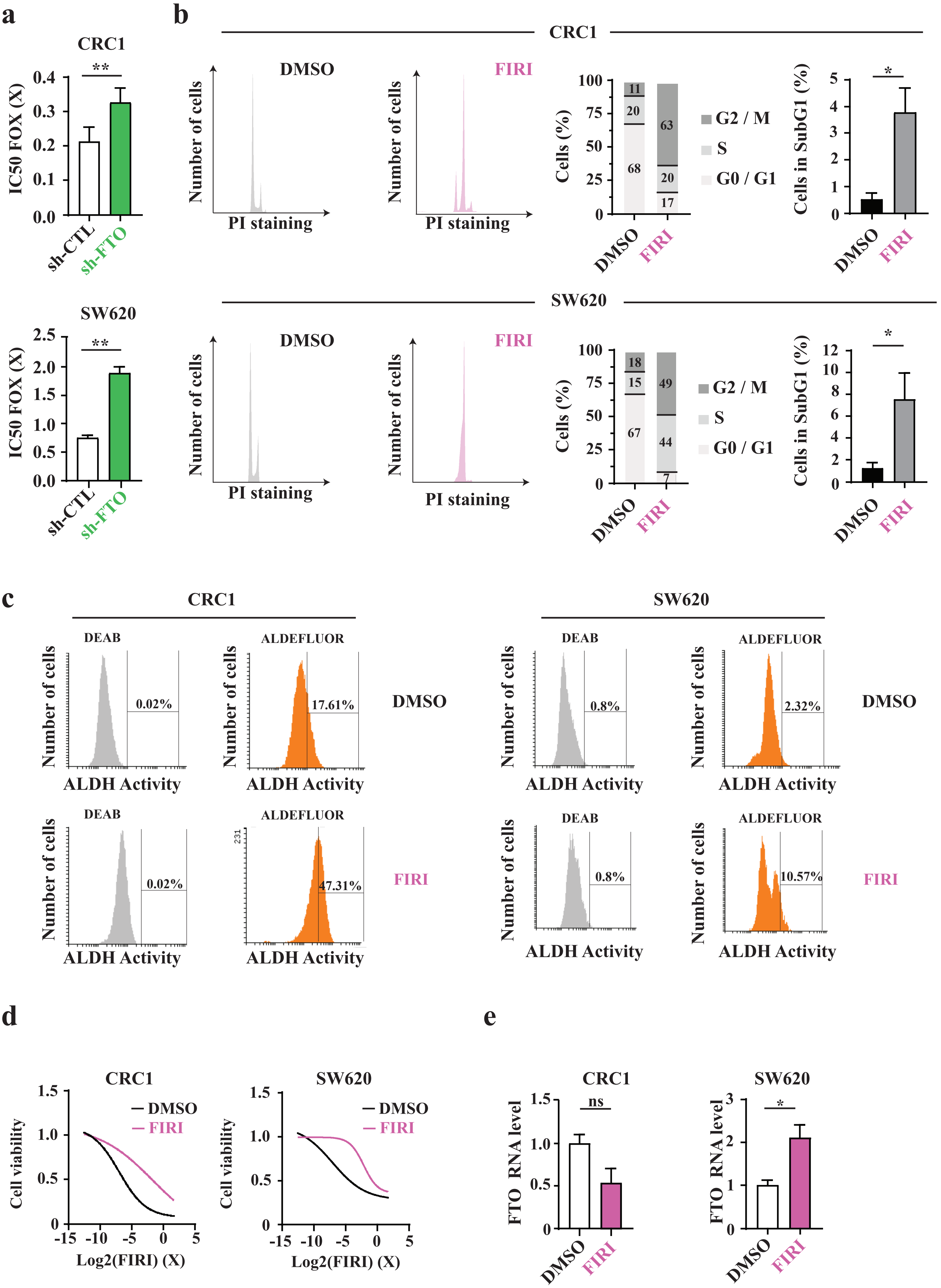

a
CRC1
IC50 FOX (X)
sh-CTL
sh-FTO
SW620
b
CRC1
DMSO
FIRI
Number of cells
PI staining
Cells (%)
G2 / M
S
G0 / G1
Cells in SubG1 (%)
SW620
c
CRC1
SW620
DEAB
ALDEFLUOR
ALDH Activity
0.02%
17.61%
47.31%
0.8%
2.32%
10.57%
DMSO
FIRI
d
CRC1
SW620
Cell viability
Log2(FIRI) (X)
DMSO
FIRI
e
CRC1
SW620
FTO RNA level
ns

**Figure S2. (a) FTO silencing increases resistance to FOX.** Cell viability was measured upon exposure of cells to increasing dose of FOX (1 X = 50 µM 5-FU and 1 µM of oxaliplatin) in CRC1 (top) and SW620 (bottom). Results represent mean +/- S.E.M of three experiments (CRC1) or two experiments representative of three (SW620). *p-value < 0.05, Two-sided Unpaired T-test. **(b) FIRI treatment blocks cell cycle in G2/M**. Cell cycle distribution was analyzed after FIRI pre-treatment at 0.2 X for 72 h in CRC1 (top) and SW620 (bottom). Cell events occurring before G0/G1 peak were considered as subG1. Bar plots represent mean +/- S.E.M of three (CRC1) and four experiments (SW620). *p-value < 0.05, Two-sided unpaired T-test. **(c) FIRI treatment enriches cell population in ALDH positive cells.** Number of ALDH positive cells after 72 h of 0.2 X FIRI treatment in CRC1 (left) and SW620 (right). Graph represents mean +/- S.E.M of three biological replicates. ** p-value < 0.01, Two-sided-Unpaired T-test. **(d) FIRI treatment selects chemoresistant cells.** Measurement of FIRI toxicity on CRC1 (left) and SW620 (right) pre-treated with 0.2 X of FIRI for 72 h**.** Cell viability was measured upon exposure of cells to increasing dose of FIRI as in 2a after a first round of 0.2X FIRI pre-treatment for 72 h. Results show one experiment representative of three. **(e) FIRI treatment impact FTO transcripts level in a cell type dependent manner**. Transcript level of FTO was analyzed by RT-PCR after 72 h of FIRI treatment at 0.2X in CRC1 (left) and SW620 (right). Bar plot represent mean +/- S.E.M of three biological replicates.

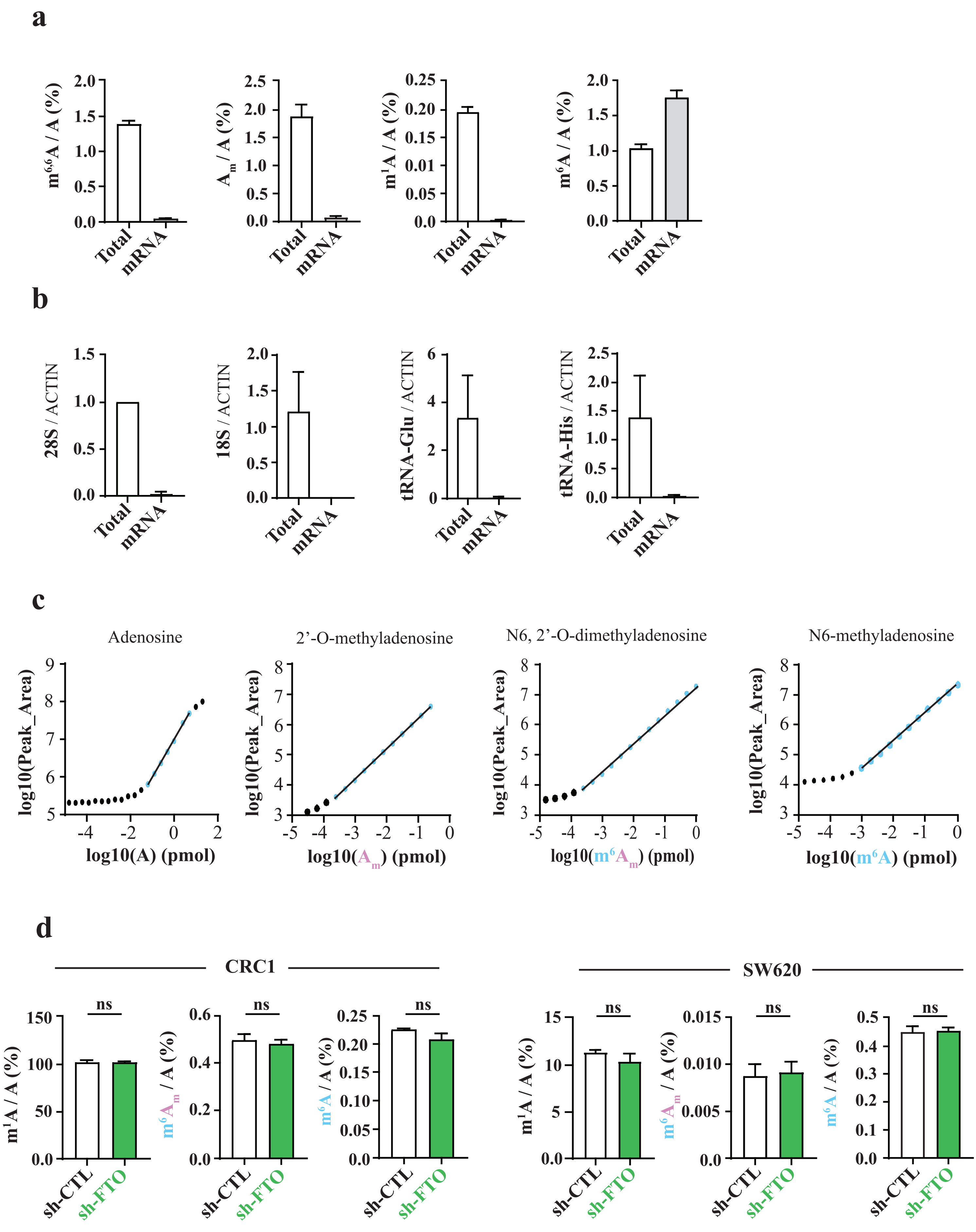

a
m6,6A / A (%)
Am / A (%)
m1A / A (%)
m6A / A (%)
Total
mRNA
b
28S / ACTIN
18S / ACTIN
tRNA-Glu / ACTIN
tRNA-His / ACTIN
Total
mRNA
c
Adenosine
2’-O-methyladenosine
N6, 2’-O-dimethyladenosine
N6-methyladenosine
log10(Peak_Area)
log10(A) (pmol)
log10(Am) (pmol)
log10(m6Am) (pmol)
log10(m6A) (pmol)
d
CRC1
SW620
ns
m1A / A (%)
m6Am / A (%)
m6A / A (%)
sh-CTL
sh-FTO

**Figure S3. (a) tRNA- and rRNA-enriched modifications sharply decrease upon mRNA purification**. Highly prevalent modifications in tRNA ($m^1A$) and rRNA ($m^{6,6}A$ ; $A_m$) or mRNA ($m^6A$) were quantified by mass-spectrometry from total RNA and polyA+ mRNA extracted from SW620 cell line to control for mRNA enrichment. Bar plots represent mean +/- S.E.M of three biological replicates. **(b) rRNA and tRNA level collapse after mRNA purification**. 28 S and 18 S rRNA, tRNA-Glu and tRNA-His levels were quantified by RT-PCR from "Total" RNA and polyA + "mRNA" extracted from SW620 cell line. Bar shows mean +/- S.E.M of 3 biological replicates. **(c) Quantifications of mRNA modifications by LC-MS/MS.** Standard curve of nucleosides standard A, $m^6A_m$, Am and $m^6A$. **(d) FTO silencing does not affect small RNA modifications.** Small RNA were purified from sh-FTO and sh-CTL cell line ( CRC1 (left) and SW620 (right)) and RNA modifications were quantified by LC-MS/MS. Bar plots represent mean +/- S.E.M of three independent experiments. ns = not significant. Two sided Unpaired T-test.

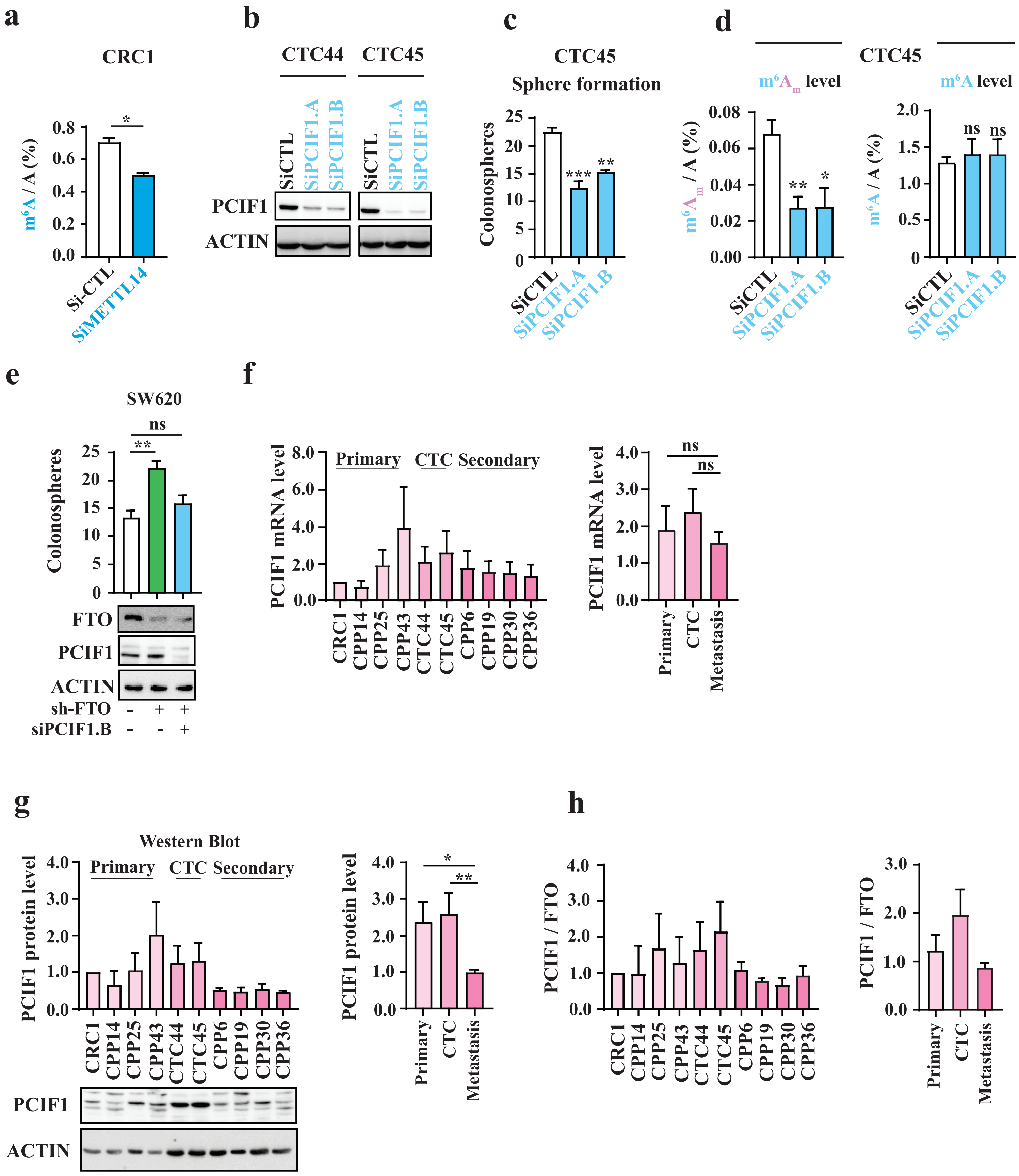

a
CRC1
m6A / A (%)
Si-CTL
SiMETTL14
b
CTC44
CTC45
SiCTL
SiPCIF1.A
SiPCIF1.B
PCIF1
ACTIN
c
CTC45
Sphere formation
Colonospheres
d
CTC45
m6Am level
m6A level
m6Am / A (%)
m6A / A (%)
e
SW620
Colonospheres
FTO
PCIF1
ACTIN
sh-FTO
siPCIF1.B
f
PCIF1 mRNA level
Primary
CTC
Secondary
CRC1
CPP14
CPP25
CPP43
CTC44
CTC45
CPP6
CPP19
CPP30
CPP36
Metastasis
g
Western Blot
PCIF1 protein level
h
PCIF1 / FTO

**Figure S4. (a) METTL14 knockdown significantly decrease m$^6$A level.** m$^6$A level of polyA + mRNA was analyzed by mass-spectrometry in CRC1 cell line. Result represents mean +/- S.E.M of two experiments. *p-value < 0.05. Two sided Unpaired T-test **(b) Control of PCIF1 depletion.** PCIF1 expression was evaluated by immunoblot upon siRNA treatment in CTCs cell lines (CTC44 (left), CTC45 (right)). **(c) PCIF1 knockdown decreases sphere formation.** Sphere formation assay was performed after siRNA treatment targeting PCIF1 in CTC45 cell line. Bar plot represent mean +/- S.E.M of at least 3 biological replicates. *** p-value < 0.001, ** p-value < 0.01. Two sided unpaired T-test **(d) PCIF1 silencing decreases m$^6$A$_m$ level in mRNA.** Quantification of polyA+ mRNA modification by mass-spectrometry upon silencing of PCIF1 in CTC45 cell line. Bar plot represents mean +/- S.E.M of 3 biological replicates. **p-value < 0.01, * p-value < 0.05 Two-sided Unpaired T-test. **(e) PCIF1 knockdown rescues basal sphere formation in sh-FTO cell line.** Sphere formation was performed upon depletion of PCIF1 by siRNA treatment of both sh-CTL and sh-FTO SW620 cell lines. Bar plot represent mean +/- S.E.M of 3 biological replicates. Proteins depletion were assessed by western blot. **p-value < 0.01, ns = not significant. One way Anova followed by multiple comparisons to sh-CTL. **(f) PCIF1 mRNA level does not change along with tumor progression.** PCIF1 mRNA level were analyzed by RT-PCR in various colon cancer cell lines that derived from primary tumor, liquid biopsies (CTC) and secondary tumor. Bar plot represent mean +/- S.E.M of PCIF1 transcript level per cell line or per group. ns = not significant, One way-Anova. **(g) PCIF1 protein level is decreased in metastatic cell line.** PCIF1 protein was quantified in patient derived cell line (same as in S5a) by western blot. Bar plot represents mean +/- S.E.M per cell line or per group. **p-value < 0.01, ns = not significant. One-way Anova followed by multiple comparisons. **(h) The ratio PCIF1 / FTO is only increased in CTCs**. Quantification of the ratio of PCIF1 protein level and FTO protein level by western blot. Bar plot represents mean +/- S.E.M of 3 biological replicates.

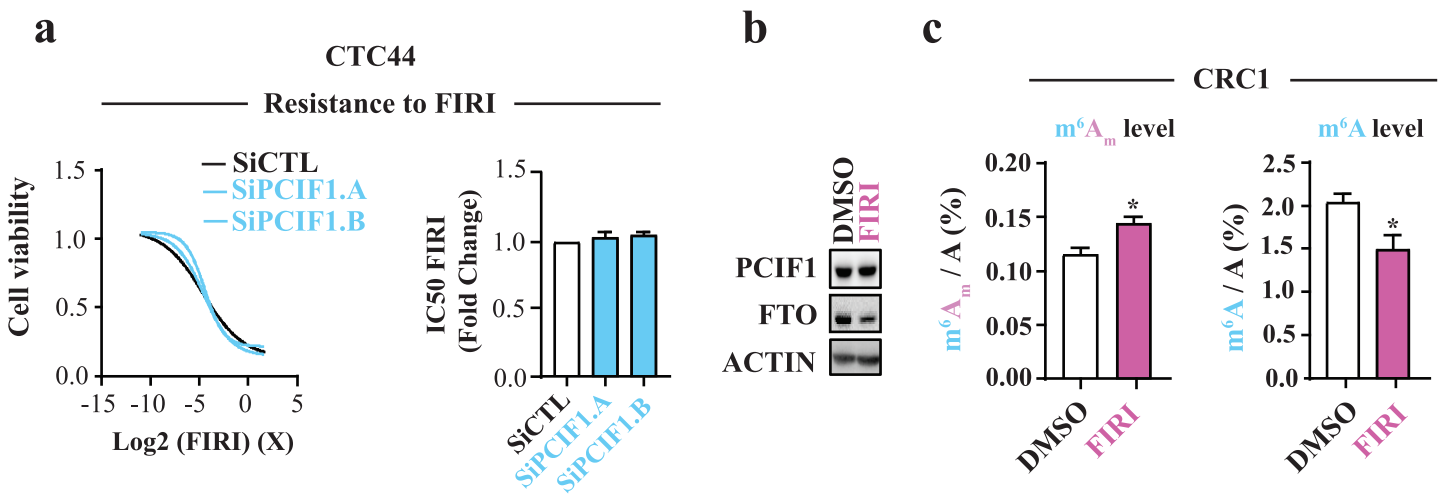


**Figure S5. (a) PCIF1 silencing does not impact chemoresistance *in vitro*.** Chemosensitivy to FIRI was assessed by treating siRNA treated CTC44 cell line with decreasing dose of drugs. IC50 was determined graphically. Bar plot represents mean +/- S.E.M of two biological replicates. **(b) PCIF1 level is unchanged upon FIRI treatment.** Analysis of FTO and PCIF1 level by western blot after cell treatment by FIRI 0.2 X for 72 h in CRC1 cell line. Picture of FTO is the same as in Figure 2d and picture of PCIF1 is representative of three biological replicates. **(c) FIRI treatment increases $m^6A_m$ level**. $m^6A_m$ / A and $m^6A$ / A level were assessed by LC-MS/MS on polyA + mRNA from CRC1 cells either treated with 0.2X of FIRI for 72 h or treated with DMSO. Bar plots represent mean +/- S.E.M of three biological replicates. *p-value < 0.05, ns = not significant, Two-sided Unpaired T-test.

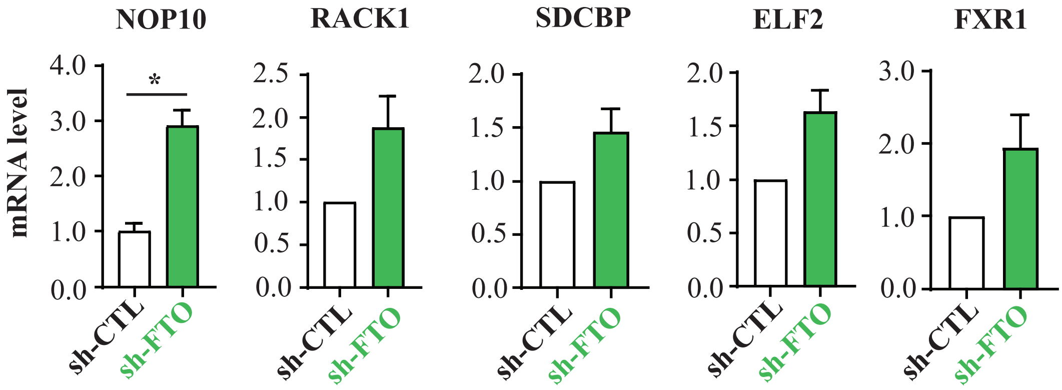


**Figure S6. Validation of differentially expressed gene in translatome data.** RT-PCR were performed on heavy fraction. Results show mean ± S.E.M of three biological replicates or two representative of three (NOP10). For each gene, cycle threshold (ct) was normalized to  ct of ACTIN. Dct were further normalized to dct of sh-CTL condition. *p-value < 0.05, Two sided Unpaired T-test.

### Supplementary Methods

#### Cell cycle analysis

Cell pellets of one million of cells were left one ice for 30 min. Then cells were resuspend in 500 µL of propidium iodide staining buffer (0.1 % of Triton X-100, 0.2 mg / mL Propidium Iodide (ThermoFisher Scientific). After one night of incubation at 4 °C in dark, cell cycle was analyzed using MACSQuant flow cytometer.

#### RNA purification

mRNA was purified from total RNA with 2 rounds of GeneElute mRNA purification kit (Sigma). rRNA was removed using Ribominus kit (Invitrogen) according to the manufacturer's instructions. Small RNA were purified from total using miRVANA kit according to the manufacturer's instructions (Invitrogen)

**Table S1: list of primers**

| Primers | Sequence 5’ – 3 ‘ |
|---|---|
| FTO-Forward | ACTTGGCTCCCTTATCTGACC |
| FTO-Reverse | TGTGCAGTGTGAGAAAGGCTT |
| ACTIN-Forward | AGCACGGCATCGTCACCAACT |
| ACTIN-Reverse | TGGCTGGGGTGTTGAAGGTCT |
| NOP10-Forward | AAACGCTTCAAGGTGCTCAT |
| NOP10-Reverse | ACCATGATTGCCTCACACAA |
| RACK1-Forward | TGAGTGTGGCCTTCTCCTCT |
| RACK1-Reverse | GCTTGCAGTTAGCCAGGTTC |
| SDCBP-Forward | TTCTGCTCCTATCCCTCACG |
| SDCBP-Reverse | CCAGTTACAGGAGCCACCAT |
| ELF2-Forward | AGGGGAATTCTTGCAAAGGT |
| ELF2-Reverse | ACACCCTTCTCTGCTCTGGA |
| PIAS1-Forward | CCTGGGTTTGTCCTGTCTGT |
| PIAS1-Reverse | AGCTCAAGCATCCATCGACT |
| 28S-Forward | CGACGTTGCTTTTTGATCCT |
| 28S-Reverse | GCAACGACAAGCCATCAGTA |
| 18S-Forward | TGTGCCGCTAGAGGTGAAATT |
| 18S-Reverse | TGGCAAATGCTTTCGCTTT |
| tRNAGlu-Forward | CCTGGTGGTCTAGTGGCTAGGA |
| tRNAGlu-Reverse | TCCCTGACCGGGAATCGAA |
| tRNAHis-Forward | GCC GTGATCGTATAGTGGT |
| tRNAHis-Reverse | TGACTCGGATTCGAACCGA |

**Table S2: list of antibodies**

| Antibodies | Manufacturers | References | Dilution |
|---|---|---|---|
| Anti-METTL3 | Abnova | H00056339-B01P | 1 : 500 |
| Anti-METTL14 | Sigma | HPA038002 | 1 : 1,000 |
| Anti-WTAP | Santa-Cruz | Sc-374280 | 1 : 300 |
| Anti-YTHDF1 | Abcam | ab99080 | 1 : 500 |
| Anti-YTHDF2 | ProteinTech | 24744-1-AP | 1 : 500 |
| Anti-ALKBH5 | Sigma | HPA007196 | 1 : 1,000 |
| Anti-FTO | Phosphosolution | 597-FTO | 1 : 1,000 |
| Anti-FTO | AbCam | EPR6895 | 1: 1, 000 |
| Anti-PCIF1 | ProteinTech | 16082-1-AP | 1 : 1,000 |
| Anti-ACTIN | Sigma | A5441 | 1 : 20,000 |
| Anti-CD44 | BD biosciences | 59942 | 1 : 500 |
| Anti-CD44v6 | Myltenyi Biotec | 130-111-238 | 1 : 300 |
| anti-IgG2a | BD biosciences | 130-091-836 | 1 : 500 |
| REA-S control Isotype | Myltenyi Biotec | 130-104-614 | 1 : 300 |
| Anti-mouse IgG | Santa-Cruz | Sc-516102 | 1 : 10,000 |
| Anti-Rabbit IgG | Cell Signaling | 7074S | 1 : 1,000 |

**Tables S3: list of si-RNAs**

| Target | si-RNA sequence 5’ – 3 ‘ |
|---|---|
| si-METTL3 | CUGCAAGUAUGUUCACUAUGA |
| si-METTL14 | AAGGAUGAGUUAAUAGCUAAA |
| si-WTAP | AAGCUUUGGAGGGCAAGUACA |
| si-YTHDF1 | CCGCGUCUAGUUGUUC AUGAA |
| si-YTHDF2 | CAGGCUGGAGAAUAACGACAA |
| si-ALKBH5 | ACAAGUACUUCUUCGGCGATT |

| | |
|---|---|
| si-FTO#1 | AAAUAGCCGCUGCUUGUGAGA |
| si-FTO#2 | ACGAAUUGCCCGAACAUUA |

**Table S4: significantly deregulated genes in transcriptome (sh-CTL vs sh-FTO)**

| ensembl_transcript_id | ensembl_gene_id | hgnc_symbol | baseMean | log2FoldChange | padj |
|---|---|---|---|---|---|
| ENST00000379284.1 | ENSG00000145990 | GFOD1 | 78,77927178 | -2,45 | 3,45E-02 |
| ENST00000502871.5 | ENSG00000138756 | BMP2K | 68,55905687 | -2,32 | 3,50E-02 |
| ENST00000560346.5 | ENSG00000137843 | PAK6 | 70,25170115 | -2,25 | 3,45E-02 |
| ENST00000613981.3 | ENSG00000276016 | ABR | 345,208247 | -1,93 | 3,45E-02 |
| ENST00000553804.5 | ENSG00000135424 | ITGA7 | 55,07126719 | -1,82 | 2,98E-02 |
| ENST00000469698.5 | ENSG00000141698 | NT5C3B | 79,73094196 | -1,71 | 3,50E-02 |
| ENST00000539396.5 | ENSG00000090372 | STRN4 | 53,85657803 | -1,63 | 3,50E-02 |
| ENST00000423729.2 | ENSG00000185818 | NAT8L | 363,546431 | -1,30 | 4,55E-02 |
| ENST00000415298.5 | ENSG00000152056 | AP1S3 | 173,4381645 | -1,23 | 2,59E-02 |
| ENST00000296978.3 | ENSG00000164484 | TMEM200A | 308,8592893 | -1,19 | 7,52E-03 |
| ENST00000268349.7 | ENSG00000140718 | FTO | 273,6964792 | 1,14 | 0,06 |
| ENST00000358075.10 | ENSG00000102158 | MAGT1 | 948,4366512 | 1,50 | 3,71E-04 |
| ENST00000540441.6 | ENSG00000166783 | MARF1 | 77,00415504 | 1,54 | 5,00E-02 |
| ENST00000453408.7 | ENSG00000141503 | MINK1 | 106,3068925 | 1,66 | 3,82E-03 |
| ENST00000635816.1 | ENSG00000107779 | BMPR1A | 236,814228 | 1,76 | 3,45E-02 |
| ENST00000369409.9 | ENSG00000092621 | PHGDH | 346,5642954 | 1,81 | 1,351E-06 |
| ENST00000508376.6 | ENSG00000134982 | APC | 316,6998041 | 1,85 | 3,28E-03 |
| ENST00000551775.5 | ENSG00000196531 | NACA | 35,05782626 | 1,86 | 3,50E-02 |
| ENST00000645393.1 | ENSG00000197467 | COL13A1 | 27,78249527 | 2,55 | 3,45E-02 |
| ENST00000620953.2 | ENSG00000274081 | PUF60 | 141,1437039 | 3,06 | 2,60E-04 |

**Table S5: significantly deregulated genes in light polysome (sh-CTL vs sh-FTO)**

| ensembl_transcript_id_gene | ensembl_gene_id | hgnc_symbol | BaseMean | log2FoldChange | padj |
|---|---|---|---|---|---|
| ENST00000473095,1 | ENSG00000106153 | CHCHD2 | 259,73024 | -4,23 | 0,00476437 |
| ENST00000573001,5 | ENSG00000006327 | TNFRSF12A | 265,6137962 | -3,77 | 0,0081455 |
| ENST00000511295,1 | ENSG00000164109 | MAD2L1 | 115,7058686 | -2,88 | 0,01551306 |
| ENST00000396296,7 | ENSG00000136758 | YME1L1 | 84,88862546 | -2,88 | 0,01228001 |
| ENST00000540297,6 | ENSG00000166598 | HSP90B1 | 109,8066838 | -2,76 | 0,04842021 |
| ENST00000546921,1 | ENSG00000170421 | KRT8 | 58,98741964 | -2,76 | 0,0028436 |
| ENST00000426663,1 | ENSG00000117475 | BLZF1 | 64,45441125 | -2,52 | 0,01087912 |
| ENST00000578758,5 | ENSG00000108654 | DDX5 | 180,4850595 | -2,42 | 0,02370278 |
| ENST00000440865,2 | ENSG00000153187 | HNRNPU | 260,0176582 | -2,28 | 0,01486871 |
| ENST00000531140,1 | ENSG00000109929 | SC5D | 96,40538527 | -2,22 | 0,03923588 |
| ENST00000583036,5 | ENSG00000265681 | RPL17 | 1353,617173 | -1,97 | 0,03056144 |
| ENST00000472729,1 | ENSG00000173163 | COMMD1 | 147,8102686 | 1,89 | 0,04127679 |
| ENST00000395577,2 | ENSG00000100526 | CDKN3 | 615,3449577 | 1,98 | 0,00476437 |
| ENST00000442660,5 | ENSG00000159147 | DONSON | 132,045702 | 2,10 | 0,01363747 |
| ENST00000614778,4 | ENSG00000249915 | PDCD6 | 536,122248 | 2,20 | 0,03899492 |
| ENST00000361166,8 | ENSG00000186575 | NF2 | 151,0018483 | 2,64 | 0,04710098 |
| ENST00000426073,6 | ENSG00000102606 | ARHGEF7 | 243,2549723 | 3,24 | 0,00688657 |
| ENST00000515539,5 | ENSG00000145781 | COMMD10 | 71,24704077 | 3,26 | 0,00983477 |
| ENST00000483423,5 | ENSG00000223766 | PRR3 | 35,70491387 | 3,52 | 0,03923588 |
| ENST00000487302,3 | ENSG00000233564 | PRR3 | 35,70491387 | 3,52 | 0,03923588 |
| ENST00000529725,1 | ENSG00000213445 | SIPA1 | 35,922234 | 3,75 | 0,0028436 |
| ENST00000582770,6 | ENSG00000105655 | ISYNA1 | 117,7625859 | 3,95 | 0,00476142 |
| ENST00000604351,5 | ENSG00000196422 | PPP1R26 | 142,5128676 | 4,39 | 1,42E-05 |

**Tables S6: significantly deregulated genes in heavy polysome (sh-CTL vs sh-FTO)**

| ensembl_transcript_id | ensembl_gene_id | hgnc_symbol | baseMean | log2FoldChange | padj |
|---|---|---|---|---|---|
| ENST00000557912,1 | ENSG00000182117 | NOP10 | 426,0909514 | -4,43 | 6,56E-14 |
| ENST00000507000,5 | ENSG00000204628 | RACK1 | 169,8339352 | -3,66 | 0,036224085 |
| ENST00000468861,5 | ENSG00000114416 | FXR1 | 334,0809707 | -2,88 | 0,028827874 |
| ENST00000522610,5 | ENSG00000070501 | POLB | 219,1002589 | -2,71 | 0,000743499 |
| ENST00000424270,6 | ENSG00000137575 | SDCBP | 255,1401181 | -2,70 | 0,03835289 |
| ENST00000577650,5 | ENSG00000141526 | SLC16A3 | 280,8281468 | -2,40 | 0,049456175 |
| ENST00000315741,5 | ENSG00000122406 | RPL5 | 1106,973304 | -2,29 | 0,028827874 |
| ENST00000373632,8 | ENSG00000118705 | RPN2 | 57,33589018 | -2,29 | 0,015093056 |
| ENST00000544233,5 | ENSG00000176871 | WSB2 | 66,42820584 | -2,28 | 0,028827874 |
| ENST00000438908,5 | ENSG00000230230 | TRIM26 | 163,5899189 | -2,16 | 0,045213002 |
| ENST00000578804,5 | ENSG00000108654 | DDX5 | 135,056588 | -2,10 | 0,036224085 |
| ENST00000319980,10 | ENSG00000032742 | IFT88 | 57,39826553 | -2,05 | 0,035151201 |
| ENST00000593027,6 | ENSG00000105677 | TMEM147 | 231,1540574 | 2,21 | 0,033499831 |
| ENST00000472481,5 | ENSG00000142541 | RPL13A | 3916,00591 | 2,22 | 0,020845154 |
| ENST00000545237,1 | ENSG00000033800 | PIAS1 | 48,03035127 | 2,31 | 0,035151201 |
| ENST00000442386,5 | ENSG00000143727 | ACP1 | 974,4346148 | 2,37 | 0,035151201 |
| ENST00000568146,1 | ENSG00000140939 | NOL3 | 119,4543559 | 2,90 | 0,048460803 |
| ENST00000394235,6 | ENSG00000109381 | ELF2 | 31,63957441 | 3,08 | 0,03835289 |
| ENST00000518015,5 | ENSG00000132912 | DCTN4 | 24,28381223 | 3,11 | 0,015093056 |
| ENST00000491937,6 | ENSG00000171863 | RPS7 | 1403,902944 | 3,20 | 0,028827874 |
| ENST00000527361,5 | ENSG00000121753 | ADGRB2 | 68,93376841 | 4,71 | 6,80E-09 |
